# Imagination as predictive perception: mental imagery predictively biases perceptual judgments of observed action kinematics

**DOI:** 10.1101/2023.06.07.544005

**Authors:** Eleonora Parrotta, Katrina L. McDonough, Patric Bach

**Author notes:** Correspondence: Eleonora Parrotta, School of Psychology, University of Aberdeen William Guild Building, King’s College Aberdeen AB24 3FX.

## Abstract

Recent approaches conceptualize mental imagery as a simulatory mode of perceptual experience, which relies on the voluntary engagement of the same top-down prediction processes that shape our perception of the external world. If so, then imagery should induce similar predictive biases as those that are known to govern the perceptual representation of others’ behaviour. In four experiments, participants saw object-directed approach and avoidance actions and reported the hands’ last seen location after their sudden disappearance. All revealed robust predictive biases, showing that perceptual judgments are illusorily distorted towards the implied goals of the actions and away from obstacles. Importantly, the experiments also showed that prior action imagery suffices to induce similar biases, so that perceptual judgments become distorted not only towards the action’s expected next steps but also the imagined ones. These imagery-induced biases were robust across stimulus sets and measurement methods. They reflect prior knowledge of how people move and can be induced not only through imagery of the actions itself, but also through imagery of situations in which the actions are merely expected. These data show that imagery induces similar perceptual expectations as other prediction processes, in line with the proposal that imagery reflects the voluntary control of predictive pathways that govern an event’s perceptual representation. Moreover, imagery can *drive* prediction processes, inducing expectations about events likely to occur in the imagined (not observed) realities, suggesting shared pathways through which imagery and prediction may support mental simulation and counterfactual reasoning.

**Public Significance Statement:** This study uses the perception of other people’s behaviour as a testing bed to advance the hypothesis that imagery can be understood as *predicted* perception: that, when people imagine, they make voluntary use of the same prediction mechanisms that otherwise allow them to anticipate – and visualise – how a situation will develop further. In four experiments, the study shows (1) that imagining another’s behaviour induces the expectation that their actions will develop in the imagined manner, (2) that imagining situations elicits expectations about how people will behave within them, (3) that these imagery-induced expectations are integrated with other expectations people have about others’ behaviour and (4) subtly distort how these behaviours are visuospatially represented. The findings demonstrate a link between imagery and predictive perceptual abilities and reveal how imagery can act as a key tool in people’s ability to anticipate relevant futures and explore counterfactual realities.

## Imagery as predicted perception: imagery predictively biases judgments of observed action kinematics

People spend much of their life imagining. In their mind, they play through future interactions with cherished (or hated) others, they anticipate future rewards, or agonize over worries that may or may not come true (Delamillieure et al., 2010; Zagacki et al., 1992). Imagery serves as a means to simulate different situations and their possible outcomes (Grush, 2004; Moulton & Kosslyn, 2009). It can serve both epistemic functions (Blajenkova et al., 2006; Finks et al., 1989) and help action planning, for when the imagined situations come to pass (Monaco et al., 2020; Mulder, 2007; Toussaint et al., 2013). Imagery and imagery-like abilities have therefore been implicated in a wide range of cognitive skills, from spatial navigation and reading comprehension to creativity, engineering, and mathematics, as well as in social abilities such as moral decision-making (e.g., Blajenkova et al., 2006; Ganis & Schendan, 2013; Pearson, 2019) and perspective taking (Ward et al., 2019, 2020, 2022).

A long-standing proposal is that imagery can fulfil these functions because it acts as a weak form of perception itself, which allows one to activate – and manipulate – the same representations as real sensory inputs (e.g., Dijkstra et al., 2017; Koenig-Robert & Pearson, 2021; Moulton & Kosslyn, 2009; Schendan & Ganis, 2012). However, while the question of whether picture-like representations underlie imagery has driven research for several decades (Kosslyn, 1973, 1980; Kosslyn et al., 1978; Kosslyn & Thompson, 2003; Pylyshyn, 1973, 1981; Shepard & Metzler, 1971), it is still unresolved. Several findings suggest close links between imagery and perception. Studies have revealed, for example, that the inspection of real and mental images follows similar time courses and elicits similar eye movements (Brandt & Stark, 1997; Laeng & Teodorescu, 2002), that imagined stimuli are represented locally in retinotopic visual and orientation space (Bergmann et al., 2016; Chang et al., 2013; Chiou et al., 2018; Naselaris et al., 2015), and that prior imagery facilitates perceptual processing of a stimulus (Crowder, 1989; Hubbard & Stoeckig, 1988) and makes it more likely that it reaches the threshold for conscious perception (Dijkstra et al., 2021; Pearson et al., 2008). Moreover, there is neuroimaging evidence that imagery activates (even early) sensory cortices (e.g., Kobayashi et al., 2004; Kosslyn & Thompson, 2003; Porro et al., 1996; Zatorre et al., 1996), and that the content of imagined scenes can be successfully decoded from fMRI activation patterns in visual cortex (Dijkstra et al., 2019; Pearson & Kosslyn, 2015).

Others have pointed out, however, that the overlaps in brain regions are much more pronounced in higher-level (i.e., frontal and parietal) regions than early sensory ones (e.g., Dijkstra et al., 2019), and that at least some of the decoding successes can be explained by re- instated eye movement patterns rather than by changes to perception itself (Mostert et al., 2018). Moreover, several other methodological confounds could give the appearance of influences on perception where there are none (see Firestone & Scholl, 2016 for a taxonomy), and it has been pointed out that even visual aftereffects induced by imagined motion (e.g., Winawer et al., 2010) can reflect decisional rather than perceptual changes (Gallagher et al., 2021). Indeed, several neuropsychological studies have revealed striking dissociations between imagery and perception (Ganis et al., 2004; Ganis & Schendan, 2008). While damage to the occipital cortex sometimes leads to impairments in visual imagery (Butter et al., 1997), even widespread damage to the early visual cortex that causes cortical blindness may leave visual imagery unaffected (Chatterjee & Southwood, 1995).

Here, we propose a subtle but important re-conceptualization of the imagery-perception-link, which treats imagery not as a form of perception itself, but as *predicted* perception. This notion dates to Neisser (1976, pp. 130–131) who argued that mental images could be understood as “a readiness to perceive the imagined object”. It recently re-emerged from predictive processing frameworks, which mirror many of Neisser’s ideas and cast perception as an iterative process of hypothesis testing and revision. On this view, perception emerges not from simple bottom-up decoding of the sensory input, but from people’s internal model of the world and their expectations of the sensory input it would generate. This internal model is kept in check by what is actually received from the senses, through a reciprocal bottom-up and top-down information exchange throughout the neuronal hierarchy, so that the model converges towards the “best guess” of the external environment (Clark, 2013; Den Ouden et al., 2012; Friston & Kiebel, 2009; Markov & Kennedy, 2013). Within such frameworks, imagery could reflect the voluntary control, and manipulation, of the brain’s internal model, by generating templates of the predicted sensory input, without being constrained by what is received from the senses (e.g., Dijkstra et al., 2019, 2020; Moulton & Kosslyn, 2009; see also, imagery as "detached schemata", Neisser, 1976). Mental imagery, in such views, would therefore not correspond to the process of *receiving* sensory input, even a weak one, but to the process of *predicting* it. In other words, it would reflect the formation of a specific hypothesis about what one currently perceives that is not kept in check by the sensory input, but which can induce perceptual expectations itself or act on the internal model from which expectations are derived.

Here, we test over four experiments whether imagery creates expectation-like effects on the processing of upcoming sensory information during motion perception. To do so, we used the classical representational momentum task, which has been designed to reveal predictive influences on the perception of moving stimuli (Freyd & Finke, 1984; for review, see Hubbard, 2005). In the task, participants simply report – via touch screen or by comparison to a probe stimulus – the perceived disappearance points of a moving stimulus. The classical observation is that these reports are predictively biased: stimuli appear to have travelled further along their (anticipated) trajectory than they really did, as if the visual system added the expected next steps to the observed motion (e.g., Hogendoorn, 2020), or filled them in after the stimulus’ sudden disappearance (e.g., Ekman et al., 2017). The resulting “motion illusions” (Hubbard, 2005) are highly robust and are reduced but not eliminated when participants are warned about their occurrence or when given feedback about the errors they make (Courtney & Hubbard, 2008; Ruppel et al., 2009). They are present for the movement of abstract dots (Hubbard & Bharucha, 1988), for objects undergoing state changes (Hafri et al., 2022), and for people moving through naturalistic scenes (Thornton & Hayes, 2004).

Importantly, the observed biases are not simple extrapolations of the observed motion paths but reflect people’s internal model about how it will develop next. They integrate multiple influences that simultaneously determine an object’s future path (e.g., Hubbard, 1994, 2005), such as how this path will be shaped by the physical forces acting upon it (e.g., gravity, Hubbard, 1995, 2020; friction, Hubbard, 1995a, 1995b; for review, see Hubbard, 2005), people’s own actions (Jordan et al., 2002), and by external cues that may predict it (e.g., verbal information whether an object will bounce off from or crash against an obstacle, Hubbard, 1994). In the social domain, the biases integrate prior knowledge of how other people’s actions typically unfold given the biomechanical constraints of the human body, as well as higher-order information about the actor’s intentions (e.g., to “take” or “leave” an object, Hudson, Bach, et al., 2018; Hudson, Nicholson, Ellis, et al., 2016; Hudson, Nicholson, Simpson, et al., 2016) and prior knowledge of how an intentional actor would most efficiently achieve these goals within the current environment (e.g., head straight towards goals but circumvent obstacles, Hudson, McDonough, et al., 2018; McDonough et al., 2019; for a preregistered replication, see McDonough & Bach, 2022).

Here, we adapt the representational momentum paradigm to test whether imagery induces such predictive perceptual distortions. If imagery acts as a form of predicted perception, then simply imagining a motion should induce similar predictive biases as those generated during motion observation itself. The observation of other people’s actions provides an ideal testing bed for this hypothesis. People are well used to imagining social situations (Honeycutt et al., 1992; Honeycutt & Gotcher, 1991; Zagacki et al., 1992, for a review see Honeycutt & Ford, 2001) and social stimuli are inherently ambiguous, making them a prime candidate for predictive influences (e.g., Bach & Schenke, 2017; Kilner et al., 2007; Tamir & Thornton, 2018). We therefore adapted the stimuli of Hudson and colleagues (Hudson, Nicholson, Ellis, et al., 2016; Hudson, Nicholson, Simpson, et al., 2016; Hudson et al., 2017), which show side views of hands in a neutral position near a potential target object, which then start to reach for or withdraw from this object. Prior to action onset, we asked participants to *imagine* one of these two actions. As soon as participants report that their mental image is vivid and clear, the action starts. It shows either the action they imagined (e.g., they imagined a reach of the object and indeed view a reach) or the respective other action (e.g., they imagine a reach but see a withdrawal). The hand disappears at an unpredictable point along the way and participants simply report the perceived disappearance point as accurately as they can, either by localizing it on a touch screen (Experiment 1), or by comparing it to a probe stimulus presented immediately after (Experiment 2).

By measuring the difference between real and reported hand disappearance points, this task allows us to reveal the multiple expectations that integrated into the prediction of the motion’s next steps (Hubbard, 2005), and test whether prior imagery forms one of these components. If so, then the data should reveal, first, the well-known predictive bias induced by motion perception itself, reflecting the expectation that moving objects will continue along their previous path (i.e., the classic representational momentum effect). People should then perceive the hands to have travelled further along the trajectory than they really did, reporting them closer to target objects than they were when viewing a reach, and further away from goal objects when seeing a withdrawal. Second, it allows us to test whether prior imagery *by itself* suffices to induce similar biases. If so, prior imagery of a reach should induce predictive biases towards the goal object while imagining a withdrawal should induce predictive biases away from it, similar to the expectations induced by actually observing these actions.

To our knowledge, this is the first test of whether imagery can induce such predictive biases in motion perception. It measures such influences, first, by using stimuli that are complex and naturalistic so that they more closely match everyday imagery processes compared to the simple abstract shapes mostly used in prior research. Second, it evaluates these biases not in terms of *whether* a stimulus is detected or identified (Pearson et al., 2008; Perky, 1910), which is particularly affected by higher-level decision biases and thresholds (Dijkstra et al., 2021), but in terms of *how* it is perceived, as illusory changes to how the stimulus is visuospatially represented. Third, the biases measured by representational momentum-like tasks reflect not the perceptual integration of two concurrent stimuli – one imagined and one observed – like previous research on imagery-perception interactions, but the combination of available sources of information into an expectation of the motion’s next steps (e.g., Hubbard, 1994; 2005). It therefore provides evidence for the proposal that imagery can induce predictive biases, which are integrated with the perceptual input similarly as expectations from actual motion (Moulton & Kosslyn, 2009).

We use the following strategy. In Experiment 1 we measure predictive biases using touch- screen judgments – when people simply report the hand’s last seen locations – and show that these judgments are biased towards both – prior motion and prior imagery – indicating an integration of both types of expectations into the internal model of the motion’s next steps.

To rule out possible contamination of results through motor or working memory biases, Experiment 2 replicates these findings by asking participants to compare the last seen stimulus to a probe stimulus presented immediately after, which shows the hand in a (future) location further along its trajectory or in a prior (past) location. Once these imagery-induced biases are established, the preregistered online Experiment 3 extends them to another stimulus set. It tests whether imagery is integrated not only with expectations derived from the observed motion, but with higher-order expectations known to shape social perception (i.e., that intentional actors head straight towards goals but will circumvent obstacles in their path). Finally, the preregistered Experiment 4 goes one step further to test whether imagery does not only integrate with higher-level expectations, but whether it can *drive* them. By asking participants to imagine not a potential action but a potential scene, we can test whether people derive expectations about how other people would achieve their goals not only from the actions available in the real environment, but from those available in the scene they currently imagine.

## Experiment 1

Experiment 1 provides a first test of whether the prior imagery of an action induces motion expectations that bias visuospatial judgments of observed action kinematics. In each trial, participants saw a hand in a neutral position near a potential goal object and imagined this hand reaching for or withdrawing from that object, depending on the object’s colour. As soon as they indicated that their mental image was vivid, (by saying “Ready!” into a microphone), the hand started to reach towards or withdraw from the object, either corresponding or not corresponding to their prior imagery. The hand disappeared at an unpredictable point in the motion sequence, and participants indicated on a touch screen the last seen location of the tip of the index finger.

The difference between the hand’s real and perceived disappearance point provides us with a quantitative measure of how people’s internal representation of the observed action differs from what is visually presented and therefore reveal how expectations about its next steps are integrated into its perceptual representation (Hubbard, 2018a, 2020). It allows us to test, first, whether participants’ judgments integrate the general expectation that the motions will continue their observed path, so that observed motions appear as more pronounced than they really were (i.e., the representational momentum effect; Freyd & Finke, 1984; Hubbard, 2005; for these stimuli, see Hudson, Nicholson, Ellis, et al., 2016; Hudson, Nicholson, Simpson, et al., 2016). Observed reaches should then be reported to have disappeared closer to an object than they really did and observed withdrawals as further away. Second, and more importantly, the study allows us to test whether imagery is integrated in people’s perceptual expectations of the action’s next steps, similarly as for other expected influences on an object’s anticipated path (Hubbard, 1994; 2005). *Imagining* a hand movement prior to their observation should then induce similar predictive biases, not towards the trajectory one expects but the trajectory one has imagined. Prior imagery of a reach should then enhance predictive biases towards the goal object but reduce those away from it. Prior imagery of a withdrawal, in contrast, should enhance predictive biases away from objects, but reduce those towards them. Such findings would provide the first indication that motion imagery induces expectation-like effects in the internal representation of observed actions, similar to expectations derived from the motion itself. To rule out that the colour cues themselves induce any biases, another (control) group of participants was not asked to imagine an action when seeing the cues, but to simply say “Ready” after they mentally counted to two. The colour of the objects should induce no biases in these participants.

### Method

#### Participants

Fifty-four participants took part in the experiment (*mean age* = 21.51, *SD* = 3.82, 39 female), recruited from the University of Plymouth and wider community. Demographic information was collected online by the recruitment system. Before the experiment, all participants were asked to declare their sex/gender. All were right-handed with normal or corrected-to-normal vision. They were randomly assigned to either the Mental Imagery group (*n* =27, *mean age* = 20.88, *SD* = 2.65, 23 female) or the No Imagery group (*n* =27, *mean age* = 22.14, *SD* = 4.68, 20 female). Seven participants, two from the Imagery Group and five from the No Imagery group were excluded due to sub-standard performance based on a priori criteria (see Exclusion Criteria). Participants gave written informed consent and received course credit for their participation. Information about the purpose of the study was provided only after the experiment was completed. The experimental procedures were approved by the Ethics Committee of the University of Plymouth.

A sensitivity analysis with G*Power 3.1(Faul et al., 2007) showed that a minimum sample size of 22 in each group provides .80 power to detect effects with Cohen’s *d*=.63. Prior studies investigating representational momentum effects with these stimuli (Hudson, et al., 2017; Hudson, Nicholson, Ellis, et al., 2016; Hudson, Nicholson, Simpson, et al., 2016) revealed that effect sizes in this experimental paradigm are consistently of this size or larger (*d* = .52 to *d* = 1.23), and piloting with the current task established that effects would be in a similar range or larger.

#### Apparatus and Stimuli

The experiment was administered using Presentation software (NeuroBS) on a Viglen DQ67SW computer and Philips Brilliance 221P3LPY monitor (resolution: 1680 X 1050, refresh rate: 60Hz). Participants wore Logitech PC120 headphones and a microphone to register verbal responses.

The stimulus set consisted of four action sequences taken from Hudson, Nicholson, Ellis, et al. (2016) and Hudson, Nicholson, Simpson, et al. (2016). They were derived from videos filmed with a Canon Legria HFS200 at 30fps and separated into individual frames with MovieDek. Stimulus sequences of 26 frames (960 X 540 px) were extracted from these sequences. They showed a right hand making natural reaches from the right side of the screen towards an object on the left (randomly selected from water glass, bottle, handle of a knife, wine glass). They showed the full reach trajectory to just before the moment of contact with the target object at approximately the 30^th^ frame of the motion sequences. Sequences were digitally manipulated using Corel Paintshop Pro X6, to replace all background details with a uniform black background. Object stimuli were modified by placing a green or red overlay of 30% opacity over them, giving them either a red or green tint, which served as the imagination cue for participants. The stimulus occupied a visual angle of 7° x 12° degrees.

The first 26 frames of each video depicting the reaching portion of the action were used for the action sequences presented in the experiment. Each action sequence could consist of 3, 4, or 5 frames (presented for 80ms each). The first frame showed the hand at a neutral middle point of the reach sequence, with a randomly selected starting frame between frames 13 to 16. The action proceeded in two frame jumps by adding 2 to the frame number for reaches (e.g., 13–15–17–19) and subtracting 2 for withdrawals (e.g., 13–11–9), respectively. The possible disappearance points therefore varied between frame numbers 17 to 24 for reaches (75 to 225 pixels distance to the edge of the goal object), and between frame numbers 12 to 5 for withdrawals (280 to 470 pixels distance to the goal object). The response stimuli were created by erasing the hand from the final frame of the sequences to depict just the object.

When displayed directly after one of the action frames, this created the impression that the hand had suddenly disappeared. Participants were instructed to report the hand’s disappearance point by touching the last seen location of the tip of the hand’s index finger.

#### Procedure

Participants completed 12 practice trials, followed by four blocks of 42 trials, resulting in a total of 168 trials for each group. Each trial (Figure 1) started with a written instruction to hold the spacebar, to ensure that participants would not track the visual stimulus with their finger. A fixation cross was then presented for 500 to 1000ms, followed by the first frame of the movie as a static image, showing the hand in a neutral position and the object either coloured red or green. In the Mental Imagery (MI) group, participants were asked to imagine the hand reaching if the object was green and withdrawing when the object was red. In the No Imagery Group, participants were asked to count to two seconds from the appearance of the first static frame. Participants were instructed to say “Ready!” into the microphone as soon as they had clearly visualised the action (Mental Imagery Group) or finished counting (No Imagery Group). This time window was calculated on the average length of time taken to imagine the hand either reaching or withdrawing in the MI group of a prior pilot group of participants. Indeed, when both groups were compared, there were no significant differences in how long both groups took to issue the “Ready!” signal, *t*(48)=.20, *p* =.83.

**Figure 1.**
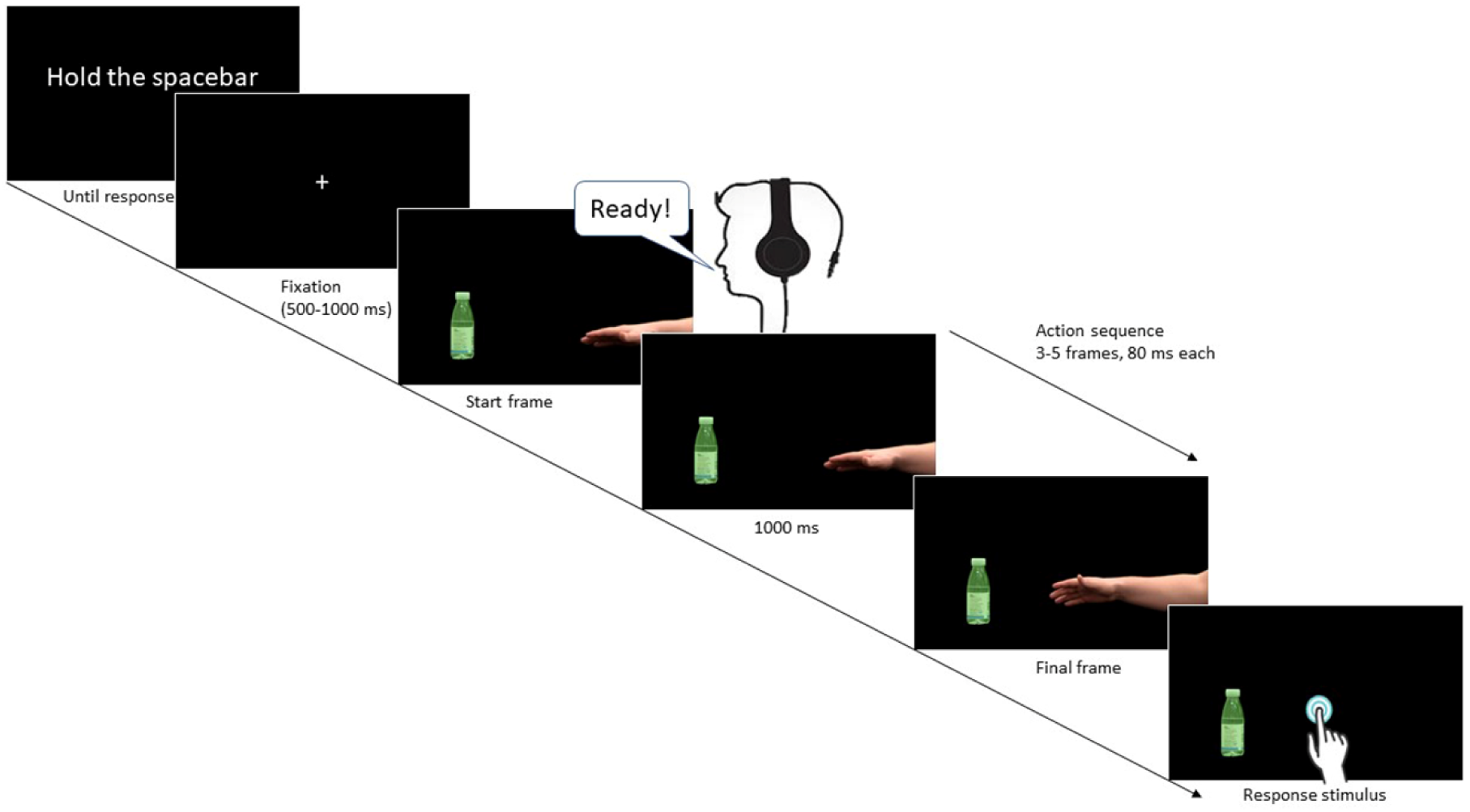
Trial sequence in Experiment 1. Participants started each trial by pressing the spacebar. They first saw a hand in neutral position near an object. Depending on colour of the object they imagined the hand to either reach for or withdraw from the object. When their imagery was vivid and clear, they said “Ready!” into the microphone, which initiated the action. After it disappeared, participants reported the hand’s index finger last seen location on a touch screen.

In both groups, the action sequence began 1000ms after the onset of the verbal response was registered by Presentation’s sound threshold logic, showing the hand reaching for or withdrawing from the object, independent of whether this appeared in red or green. After the hand had disappeared mid-action after 3, 4, or 5 frames (presented for 80ms each), participants released the spacebar and located the last seen position of the tip of the index finger on the touch screen.

After the consent and instruction phases and before the practice and experimental trials were administered, participants were asked to passively watch a demo of the action sequences they would later view and imagine (12 trials). Action sequences shown in this phase were the same as those presented in the experiment, with the only difference being that the sequence length was fixed (5 frames). The object colour was consistently allocated to the relevant action direction, for both the Mental Imagery and the No Imagery group, as it was in the following experimental phase (i.e., green = reach; red = withdrawal). This allowed participants to familiarize themselves with the stimuli for the subsequent imagery task and helped them form robust associations between object colours and the action they would have to visualize afterward. The assignment of colours to actions was not counterbalanced across participants, in order to alleviate retrieval needs that would arise from more arbitrary colour- action associations and to avoid any source of participant confusion during the imagery task.

#### Questionnaires

After the experiment, all participants completed the Vividness of Visual Imagery Scale (VVIQ; Marks, 1973), which is a 16-item self-administered questionnaire that assesses the subjective vividness of visual imagery. The VVIQ was chosen over the more recent VVIQ2 (Marks, 1995), due to its shorter length but roughly equivalent psychometric properties (Campos & Pérez-Fabello, 2009). Each item asked participants to form a specific mental image and then rate this image on a five-point scale, ranging from 1, “No image at all”, to 5, “Perfectly clear and vivid as normal vision”. Traditionally, participants make 16 ratings with their eyes open, then 16 ratings with their eyes closed, and the two sets of scores are added together. However, McKelvie’s (1995) meta-analytical review found no difference in the ratings made with eyes open and eyes closed. Thus, the participants only completed the questionnaire once with their eyes open.

Additionally, participants in the Mental Imagery group filled out introspective reports about their imagery experiences (Jack & Roepstorff, 2002). The introspective report asked for a written description of the imagery experienced. It was followed by a specific question about their use of *kinaesthetic* (defined as a ‘feeling or physical sensation’) and *visual* imagery strategies, which they rated, on a Likert five-point scale, the intensity of the muscular sensations and the clarity of the visual images they experienced. This was followed by a question asking which of these strategies subjects personally relied on during the experiment (visual only, kinaesthetic only, or combined visual and kinaesthetic).

Research has also highlighted that the *capacity to control* the action occurring in an image, as well as the clarity and *vividness* of the image as important dimensions (Goss et al., 1986; Housner & Hoffman, 1981; Isaac & Marks, 1994). Therefore, we explored whether individual differences along this ability could also elicit a corresponding variation in shaping action visual perception. Participants rated on a Likert five-point scale how much they agreed/disagreed with two statements regarding the vividness (“*The mental image was clear and vivid*”) and the controllability (“*The mental image was easy to control*”) of the experienced mental imagery, for both action directions (i.e., reaches and withdrawals).

#### Exclusion criteria

The same exclusion criteria were applied in both groups, based on prior work with this experimental paradigm (Hudson, Bach, et al., 2018). Trials (2.7%) were excluded when the response time was faster than 200ms, or 3*SD* slower than the mean across groups, or when the selected position of hand disappearance was more than 200 pixels away from its real disappearance point. Participants were excluded (Mental imagery group, *n* = 2, No imagery group, *n* = 5) if their responses showed no consistent relationship with the visual stimuli, indicating a lack of task engagement (see Hudson, Bach, et al., 2018), based on the following criteria: (1) if the mean screen distance (pixels) between the real and selected screen positions was more than 3*SD* above the group mean, (2) if the correlation coefficient between the real and selected screen positions on both axes was smaller than .80, (3) they had less than 70% valid trials post exclusion (no exclusions).

### Results

Analysis followed our previous work with this procedure (Hudson et al., 2017). To provide a measure of each participant’s bias in localization responses, the real final screen coordinate of the tip of the index finger was subtracted from participants’ selected screen coordinate on each trial. Analysis was conducted on this localization error, which provided a directional measure of how far, in pixels (px), participants’ localisation judgments were displaced along the X- and Y-axes. An accurate response would produce a value of 0 on both axes. On the X-axis, positive values denote a rightward displacement away from the object, and negative values a leftward displacement towards the object. On the Y-axis, positive and negative values denote upward and downward displacements, respectively.

Each participant’s average differences values were entered into a 2 X 2 X 2 mixed measures analysis of variance (ANOVA) for the X- and Y- axes separately, with Group (Mental Imagery vs. No Imagery) as a between-subjects factor, and Imagery Prompt (Red vs. Green) and Action (Reach vs. Withdrawal) as within-subjects factors (Figure 2). Main effects of Action will reveal the extent to which prior action kinematics (reaches vs. withdrawals) bias perceptual reports in the prior motion direction (further towards the object for reaches, further away for withdrawals) and will therefore indicate the presence of the representational momentum effect (Freyd & Finke, 1984a; Hubbard, 2005). Note that touch screen judgments are affected by various biases, such as a bias towards the target’s centre of gravity. Main effects of Imagery Prompt will reveal the extent to which responses are biased by mental imagery in the Mental Imagery group and by the mere association between the colour and action direction in the No Imagery group. The prediction is that perceptual judgments in the Mental Imagery group would be displaced towards the imagined trajectory, that is, leftwards when reaches are imagined for green objects and rightwards when withdrawals are imagined for red objects [Footnote ^1^]. If these biases are caused by the preceding imagery task and not simply the association of colours to actions, we expected this effect to be reduced in the No Imagery group. Interactions with Group will therefore reveal whether groups differ in how much perceptual judgments are biased by either kinematics (Action X Group) or by the presence/absence of mental imagery processes (Imagery Prompt X Group). As we have no further predictions, all other main effects and interactions should be treated as incidental unless meeting a (Bonferroni-adjusted) alpha threshold of .008 to account for hidden multiplicity in an ANOVA (Cramer et al., 2016).

**Figure 2.**
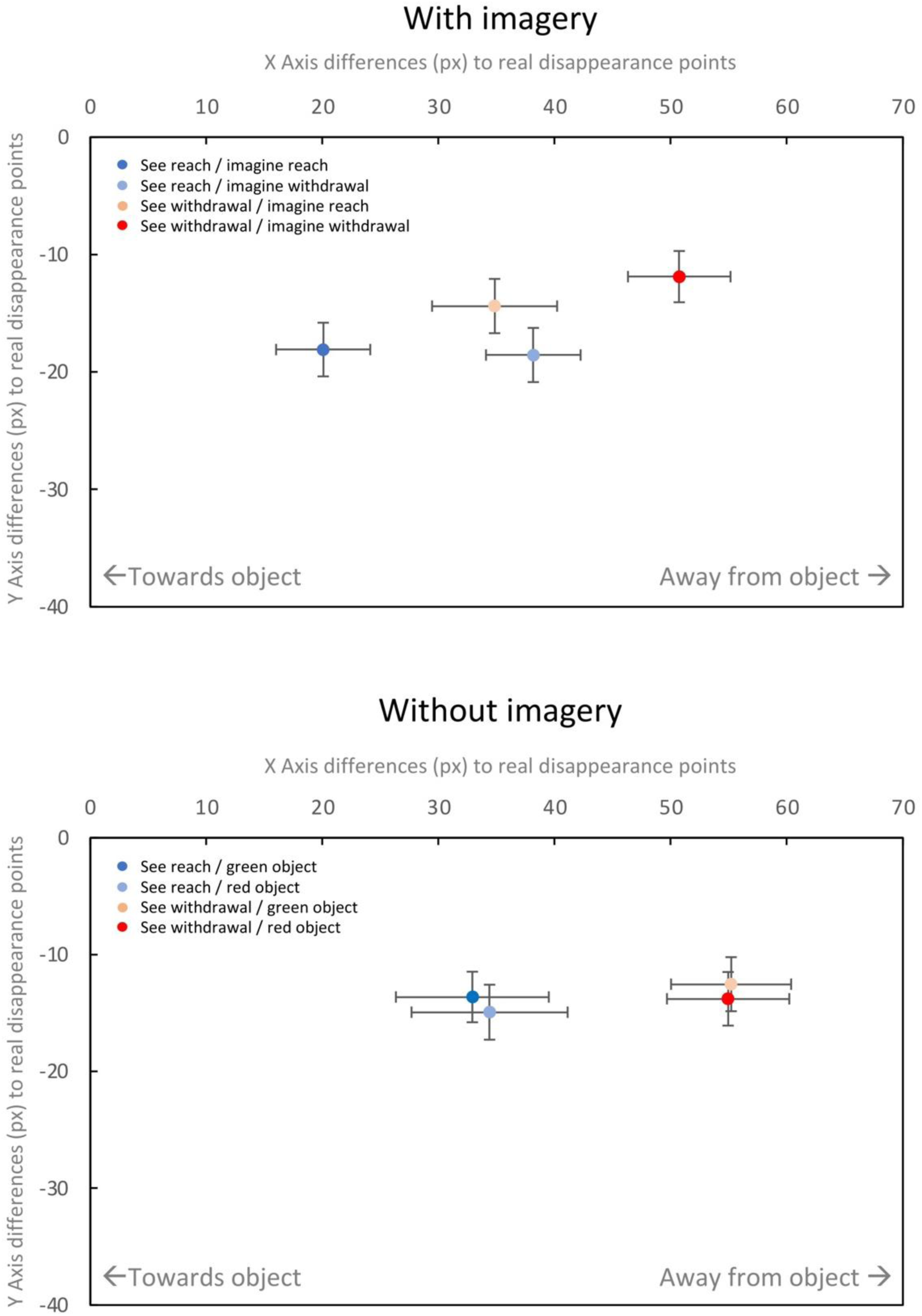
Differences on the X and Y axis between participants’ reported hand disappearance points relative to their real disappearance points (0, 0). On the X-axis, positive values denote a rightward displacement away from the object, and negative values a leftward displacement towards the object. On the Y-axis, positive and negative values denote upward and downward displacements, respectively. Accurate responses would produce a value of zero on both axes, as zero coordinates represent the hand’s vanishing point. Error bars show the standard error of the mean of the main effect of Imagery Prompt in each group (Loftus & Masson, 1994).

**Figure 3.**
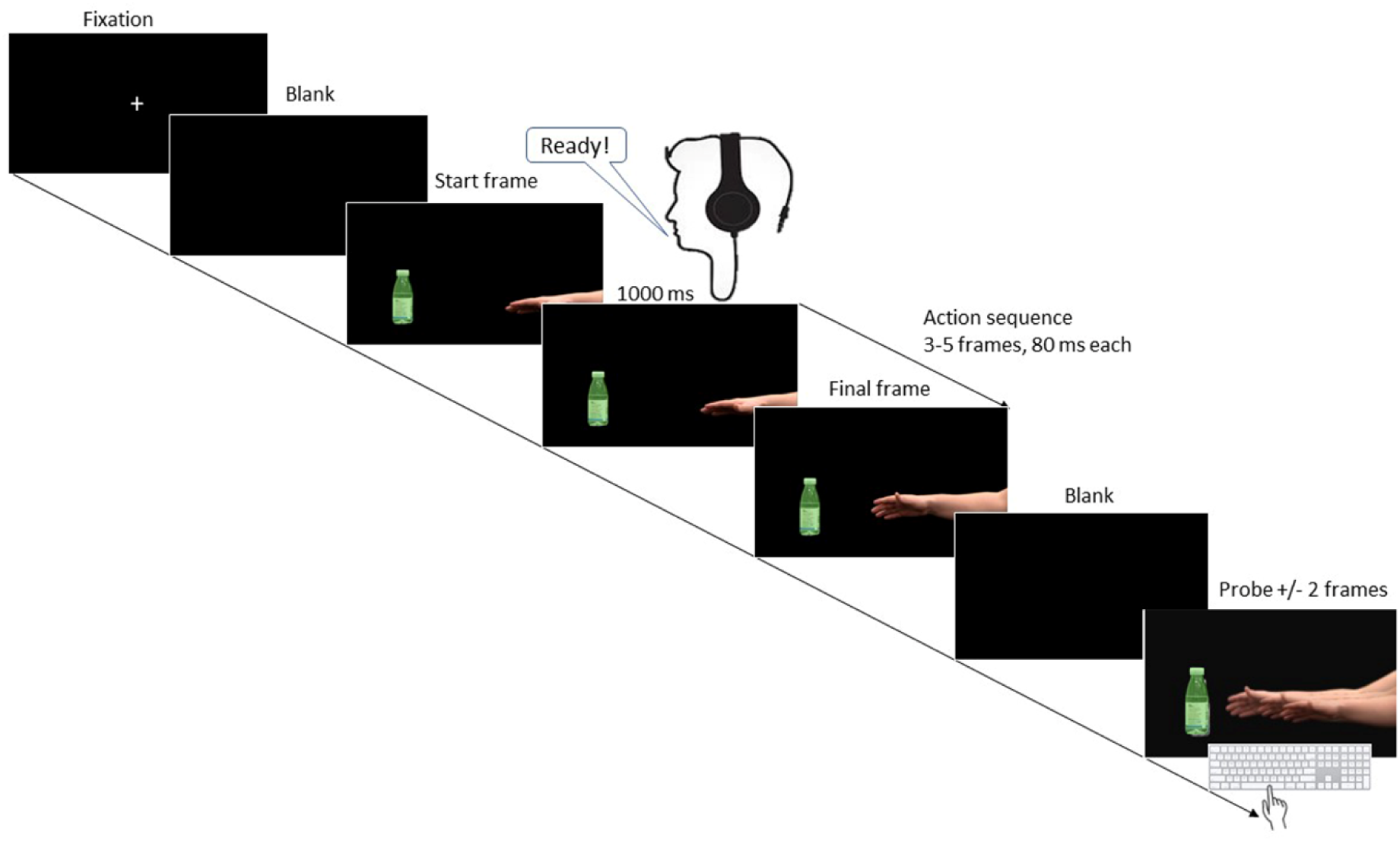
Schematic of the trial sequence in Experiment 2. Participants first saw a hand in neutral position near an object. Depending on colour of the object they imagined the hand to either reach for or withdraw from the object. When their imagery was vivid and clear, they said “Ready!” into the microphone, which initiated the action. After it disappeared for 250ms, the probe stimulus appeared, showing the hand in the same position, 1 or 2 steps further towards the object, or 1 or 2 steps further away from the object. Participants pressed the space bar when this probe stimulus was different from the last seen position of the hand.

#### X-axis

Figure 2 shows the influence of Imagery Prompt and Action on the size of the perceptual bias for both groups. Consistent with prior work with touch screen responses (e.g., Coren & Hoenig, 1972; Hudson, McDonough, et al., 2018; McDonough et al., 2019), responses were generally biased rightward (40 px) and downward (-14 px) across both groups, towards the hand’s centre of gravity. Due to these general biases affecting location responses, it is not possible to estimate the presence of representational momentum against an objective zero point (but see Experiment 2 for such a test). Instead, as in our prior research (Hudson et al., 2017), it is indexed through the *difference* in perceptual biases for reaches and withdrawals: whether seeing reaches biases perceptual errors more strongly towards the object than seeing withdrawals, which should bias them more strongly away from the objects. As expected, the ANOVA revealed the relevant main effect of Action, *F*(1,45) = 16.8, *p* < .001, *ηp^2^* = .27.

Participants’ localisation errors were biased more leftwards (i.e., towards the object) after seeing left-ward going reaches than after seeing rightwards-going withdrawals, confirming the expected displacement in the direction of motion. Action did not interact with Group, *F* < 1, consistent with the notion that this basic predictive extrapolation of motion is derived from the stimulus itself and is independent of whether people imagine an action beforehand or merely count to two.

Importantly, as expected by the proposal that imagery can induce similar predictive biases as the expectations derived from actual motion perception, the ANOVA revealed a main effect of Imagery Prompt (*F*(1,45) = 34.0, *p* < .001, *ηp^2^* = .445), showing that green objects/imagined reaches generally produce a larger leftward displacement than red objects/imagined withdrawals, even though identical action sequences were actually observed. Importantly, and as predicted, this main effect was qualified by an interaction of Imagery Prompt and Group, *F*(1,45) = 34.9, *p*<.001, *ηp^2^* = .437. The colour of the object affected the perceptual bias more strongly in the Mental Imagery group than in the No Imagery group. Indeed, step-down analyses revealed an effect of Imagery Prompt on the perceptual bias in the Mental Imagery group, *F*(1,24) = 44.5, *p* < .001, *ηp^2^*=.65) but not in the No Imagery Group, *F* < 1.

#### Y-axis

We did not have predictions about how imagery processes would affect displacements on the Y-axis, so all findings should be interpreted with caution unless they surpass a (Bonferroni) adjusted threshold for multiple comparisons in exploratory ANOVA of *p* < .008 (Cramer et al., 2016). The analysis revealed no main effect of Action, *F*(1,45) = 4.63, *p* = .037, *ηp^2^* = .09, no main effect of Imagery Prompt, *F* < 1, and no interaction, *F*(1,45) = 5.55, *p* = .023, *ηp^2^* = .110. No interactions with group passed the corrected threshold, all *F*(1,45) < 4.69, *p* > .036, *ηp^2^* < .094.

### Discussion

Experiment 1 replicated the classic predictive perceptual bias in motion perception, showing that people perceptually extend motions they have seen towards how they expect them to continue (i.e., Freyd & Finke, 1984; Hubbard, 2005; Hudson, Bach, et al., 2018; Hudson, Nicholson, Ellis, et al., 2016; Hudson, Nicholson, Simpson, et al., 2016). Hands reaching for objects were therefore reported to have disappeared closer to the objects than they really did, and hands withdrawing from them were reported to have disappeared further away. These findings are consistent with the idea that the brain represents motion in terms of its expected next steps, and “fills in” these next steps when the stimulus suddenly disappears (Ekman et al., 2017). The resulting bias in perceptual judgments was present both when participants imagined actions before they saw the stimuli, and when they merely counted to two, in line with a largely automatic visual bias that is derived from the visual stimulus (Freyd & Finke, 1984; Hubbard, 2005).

Importantly, the study showed for the first time that imagery was able to induce similar expectations as those derived from seeing the motion itself. Imagining a reach towards the object caused participants to report the same moving hands to have disappeared closer to this object, while imagining a withdrawal biased judgements away from the objects. Our findings therefore provide a first demonstration that imagery induces similar motion expectations as the observation of motion itself and that, during motion perception, both sources jointly shape perceptual judgments, in line with the Bayesian-like integration of different influences into expectations of the action’s next steps (Hubbard, 1994, 2005, 2020). Crucially, this effect of imagery was only found when participants actively imagined the two types of actions; it had no such effects when participants saw the same action stimuli and associated colour cues but did not imagine movements of the hand in either direction. This absence of the perceptual distortions in this group rules out that the perceptual bias emerged from the association between the action and the colour of the object and underlines the causal role of imagery processes in inducing motion expectations.

## Experiment 2

Experiment 1 provided the first evidence that visual imagery induces expectation-like biases, which become integrated with motion expectations generated during the perception of motion itself. However, the touch screen judgments in Experiment 1 leave open whether these estimations only reflect the perceptual representations of the observed actions, or whether they may also reflect changes in the action’s representations in working memory or on the motor level, for example, in the sensorimotor maps that guide the finger movements to the relevant locations on the touch screen (Kerzel, 2003; Müsseler et al., 2008). Experiment 2 therefore replicated the imagery effect of Experiment 1 with a psychophysical probe detection task that is unaffected by such motor or working memory influences. In each trial, participants compared the hand’s last seen position to a probe stimulus that was presented directly after hand offset (250ms gap), which showed the hand either in a future position on its trajectory than it really disappeared (e.g., closer towards the object when seeing a reach, further away for withdrawals) or on a past position (e.g., further away from the object than it really was when seeing a reach, closer for withdrawals). They simply pressed a button when they judged the probe hand’s position to be different from the hand’s last seen position on the screen, providing a direct measure of the changes to their perceptual representation.

As in Experiment 1, we predicted, first, that probe stimuli shifted in the direction of the seen motion (e.g., towards the object for reaches and away from the object for withdrawals) should be more likely to be mistaken for the hand’s last seen position, reflecting the well-known representational momentum effect (e.g., Freyd & Finke, 1984; Hubbard, 2005). Second, if imagery induces motion expectations, then participants should be more likely to mistake probes shifted further ahead in the imagined direction with the hand’s last seen position, compared to shifts against the imagined direction. Importantly, the probe judgment task makes it possible to estimate the amount of representational momentum for each action separately against the hand’s actual disappearance point, which was not possible with the touch screen task of Experiment 2. Under the assumption that the measured expectations towards the motion’s next steps combine multiple influences (Hubbard, 1994, 2005), participants should therefore be most likely to mis-identify probes further ahead in the direction of motion with the hand’s actual disappearance point when they saw and imagined the same action and both influences could summate, compared to when participants saw and imagined different actions, where both influences would cancel each other out.

### Method

#### Participants

Thirty-four participants took part in the experiment (*mean age* = 23.10, *SD* = 7.35, 23 female). Subjects were recruited from the University of Plymouth and wider community. They gave written informed consent and received course credit for their participation. Eight participants were excluded due to performance (see Exclusion Criteria). All participants were right-handed with normal or corrected-to-normal vision. Information about the purpose of the study was provided only after the experimental tests were completed. The experimental procedures were approved by the Ethical Committee of the University of Plymouth (United Kingdom). Sensitivity analysis with G*Power 3.1 (Faul et al., 2007) revealed that a sample size of 28 provides .80 power to detect effects with Cohen’s *d* = .55.

#### Apparatus, stimuli and procedure

The apparatus and stimulus set used for Experiment 2 was the same as in Experiment 1. The only difference consisted in the absence of the touchscreen for recording participants’ localization responses. Instead, subjects judged the hand’s disappearance point relative to probes that were either identical to the hand’s last seen position or were shifted along its trajectory, either towards or away from the object.

In the experiment proper, each trial began with a fixation cross followed by a blank screen (combined duration 1000ms), followed by the first static frame of the action sequence, showing the hand in a neutral position and the object, either coloured red or green.

Participants were required to imagine the hand reaching towards the object if the object was green and withdrawing from the object when the object was red. Subjects were then instructed to say “Ready!” into the microphone as soon as they felt they had clearly visualised the action. The action sequence was triggered 1000ms after the onset of the verbal response was registered by Presentation’s sound threshold logic, showing the hand reaching for or withdrawing from the object, independent of whether it appeared in red or green.

After the hand had disappeared mid-action after 3, 4, or 5 frames (80ms. each), a blank screen was presented (250ms), followed by a probe stimulus (4000ms or until response). This consisted in a single frame taken from the same action sequence. It showed the hand in either the same position as the final frame of the action (“0”) or showing a step further along the reach (i.e., “+” shifted towards the object), or not as far along the reach (i.e., “-” shifted away from the object). The probes were shifted either 1 frame or 2 frames further towards or away from the objects, producing 5 possible probe positions (2, 1, 0, +1, +2). In addition, probes of 4 frames further toward or away from the object (+4, -4) were included as rare catch trials to identify participants who were insensitive to even these largest probe distances but were not analysed (see below). Participants pressed the spacebar if they thought that the probe position was different from the hand’s final position and did not respond if they thought that it was the same.

Participants completed the exact same familiarisation trials as in Experiment 1 (i.e., demo of action sequences), to aid their imagination of the two types of actions. They then completed 12 practice trials that were identical to the main experiments, followed by four blocks of 42 trials each, resulting in a total of 168 trials. Each of these trials had an 8% chance of being a catch trial, in which the probe was shifted by 4 frames from the final position of the action sequence. As in previous studies (Hudson, Nicholson, Ellis, et al., 2016; Hudson, Nicholson,

Simpson, et al., 2016), these catch trials were used to identify participants who did not comply with task instructions and did not distinguish even the largest differences between probe and actual disappearance points (see below: Exclusion Criteria) but were not analysed. No control group without imagery instructions was tested in Experiment 2.

#### Exclusion Criteria

Exclusion criteria were based on prior work with this experimental paradigm (Hudson, Nicholson, Ellis, et al., 2016; Hudson, Nicholson, Simpson, et al., 2016). Participants were excluded if they failed to identify a larger proportion of probes as different from the real disappearance points when the probes were shifted by ±4 frames in or against the direction of motion in the catch trials, compared to the subtler shifts (±1 or ±2 frames) in the experimental trials (*n* = 5). Three additional participants were excluded because they misunderstood the task, responding not when the probe differed from the real disappearance points, but when the observed action differed from imagined one, as indicated by a larger than 90% difference in key presses for trials in which observation and imagery matched than when they mismatched.

### Results

#### Main analysis

To measure how much participants’ identification responses were affected by the action they saw and the action they imagined, we entered the proportions of individual “different” responses across probe distances into a repeated measures ANOVA with the within-subject variables Imagery Prompt (Green vs. Red), Action (reach vs. withdrawal), Probe Direction (towards the object, away from the object), and Probe Distance (2 frames vs. 1 frame) as repeated measures factors. Interpretation follows prior work (i.e., Hudson, Nicholson, Ellis, et al., 2016; Hudson, Nicholson, Simpson, et al., 2016). Main effects of Probe Distance will reveal whether participants are better at identifying probes shifted further from the real disappearance points (i.e., 2 frames forward or backward) compared to probes shifted not as far (1 frame forward or backward) and therefore indicate appropriate task engagement.

The main experimental predictions are tested by the two 2x2 interactions. First, the interaction of Action and Probe Direction will reveal to what extent perceptual judgements are biased by the action that was observed, biasing responses further towards the object when reaches were observed, and away from the object when withdrawals are observed. It is mathematically equivalent to the comparison of whether people are more likely to accept probes shifted towards the object than away from it after seeing a reach compared to seeing a withdrawal. Interactions of Imagery Prompt and Probe Direction will reveal whether perceptual judgements are similarly biased by the action that was *imagined* prior to action onset, biasing responses further towards the object when reaches were imagined, and away from the object when withdrawals were imagined. It is mathematically equivalent to the test whether people are more likely to accept probes shifted towards the object than away from it after *imagining* a reach compared to *imagining* a withdrawal.

The results are shown in Figure 4. As expected, the ANOVA showed a main effect of Probe Distance, *F*(1,25) = 71.9, *p* < .001, *ηp^2^*= .742, with larger differences between probe frames and real disappearance points (±2) detected more readily than smaller differences (±1). The predicted two-way interaction of Action and Probe Direction, *F*(1,25) = 12.0, *p* = .002, *ηp^2^*= .324, showed the expected predictive perceptual bias in the direction of motion (i.e., representational momentum). Participants accepted more readily as “same” probes that were shifted in the direction of motion, meaning more “same” responses for probes shifted towards the object when viewing a reach and more “same” responses for probes shifted further away from the object when viewing a withdrawal. Finally and most importantly, the ANOVA showed the predicted interaction of Imagery Prompt and Probe Direction, *F*(1,25) =18.3, *p* < .001, *ηp^2^* = .423, revealing that imagery of reaches induces similar predictive biases as the observation of motion itself, as seen in Experiment 1. After imagining a reach, participants were more ready to accept as “same” probes shifted towards the object (compared to away from it). In contrast, after imagining a withdrawal, participants were readier to accept as “same” probes shifted further away from the object (compared to towards it). There were no other main effects or interactions, *F* < 2.7, *p* > .11 for all.

**Figure 4.**
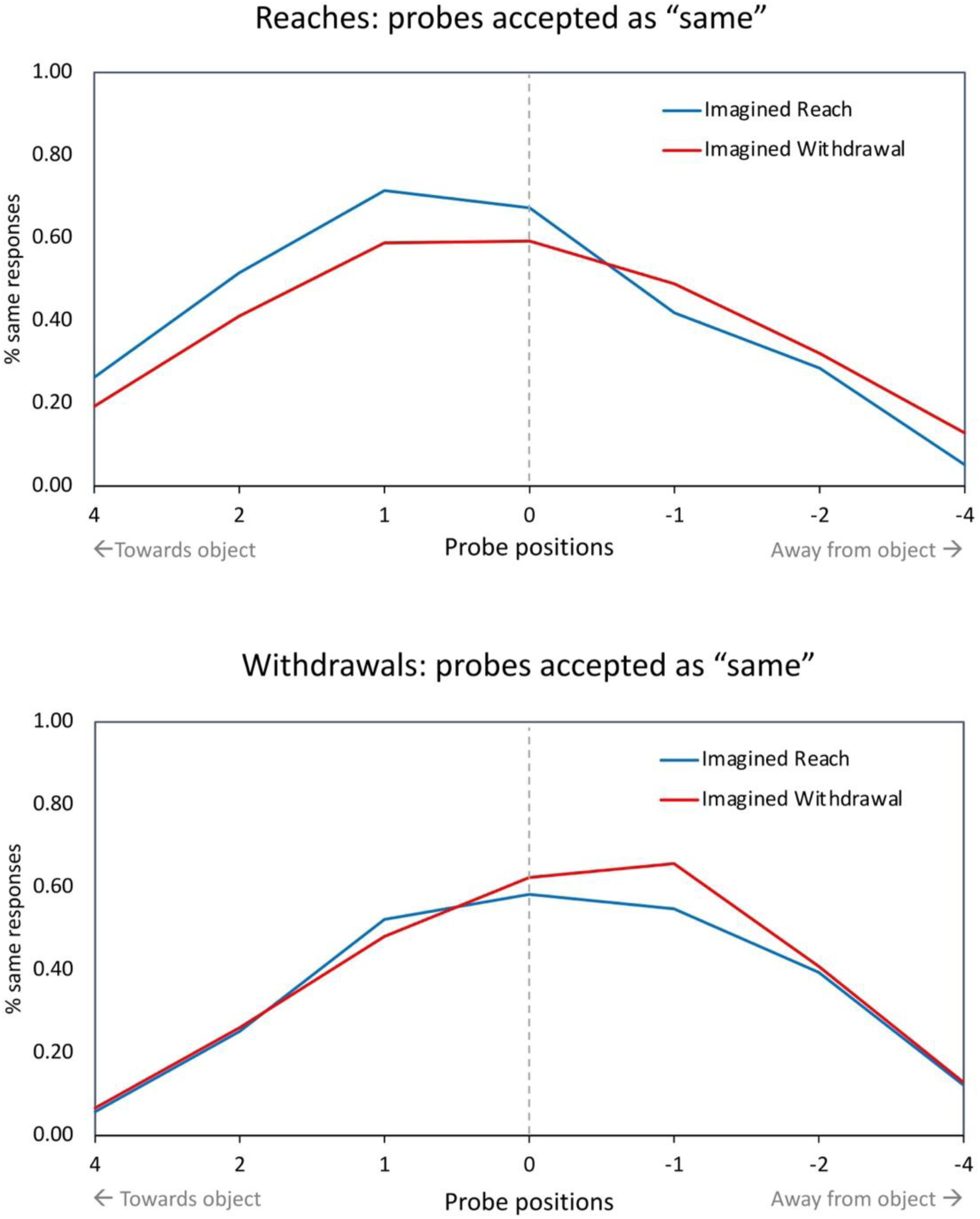
Proportion of same responses for each of the different probe stimuli, depending on whether and how far the probe was shifted further along the direction towards the object (positive numbers), was identical with the last seen position (zero), or further away from the object (negative numbers), plotted separately for observed reaches (top panel) and observed withdrawals (bottom panel), and when participants imagined reaches (blue lines) or imagined withdrawals (red lines). Same probes (0) and probes with maximal distances (±4 steps, catch trials) are shown for illustrative purposes but were not analysed.

Step-down analyses of each condition separately showed that representational momentum, calculated of increased likelihood of accepting probes shifted in the direction of motion as “same” compared to probes shifted against the direction of motion, was present in the conditions when imagery and direction of motion matched (imagining and seeing a reach, *t* = 4.85, *p* < .001, *d* = .95; imagining and seeing a withdrawal, *t* = 2.30, *p* = .030, *d* = .45), but not when they mismatched (imagining a reach but seeing a withdrawal, *t* = 1.37, *p* = .183, *d* = .27; imagining a withdrawal and seeing a reach, *t* = 1.60, *p* = .123, *d* = .313). This is consistent with the predicted additive integration of expectations generated from the seen motion kinematics and prior imagery, which summate when congruent but cancel each other out when incongruent (cf. Hubbard, 1994; 2005).

#### Additional analysis: weighted means

In the prior literature (Freyd & Jones, 1994; Hubbard, 1993; Munger et al., 1999; Ruppel et al., 2009), the results of representational momentum tasks are often analysed in terms of the weighted means, in which representational momentum for each participant in each condition is calculated as the sum of the products of the proportions of same responses and probe distances relative to the last seen image (+2, +1, 0, -1, -2 frames shifted towards or away from the object), divided by the sum of the proportions of same responses across probe distances. This analysis treats each “same” response as a point in a coordinate system, so that more distant probes identified as “same” have a larger contribution than those closer to the last seen frame. As in the main analysis, zero values indicate the absence of perceptual errors.

Positive values indicate biases in mis-localisations of hand disappearance points towards the object and negative values reflect shifts away from it.

The analysis replicates all relevant results of the main analysis. As Probe Distance and Probe Direction are already incorporated into the calculation of the weighted mean, only the factors Action and Imagery prompt are included in the ANOVA. The main effect of Action captures the representational momentum effect and revealed robust differences, *F*(1,25) = 13.2, *p* < .001, *ηp^2^*= .346. Indeed, participants were more likely to accept as “same” probes shifted towards the object compared to away from it after seeing reaches than seeing withdrawals. The main effect of Imagery Prompt captures that Imagery induces similar predictive biases in perceptual judgments towards or away from the object, *F*(1,25) = 9.45, *p* = .005, *ηp^2^*= .274. As in the main analysis, participants were therefore more likely to accept as “same” probes shifted towards the object after merely *imagining* reaches than after *imagining* withdrawals.

Follow-up analyses tested whether representational momentum could be observed in each condition separately (often termed “M-displacement” in previous work on representational momentum, Hubbard, 2005; Hubbard & Bharucha, 1988). This analysis confirmed robust displacements in the direction of motion when imagery and direction of motion were congruent (imagining and seeing a reach, *M* = .36, *SD* = .40, *t* = 4.54, *p* < .001, *d* = .89; imagining and seeing a withdrawal, *M* = -.21, *SD* = .49, *t* = 2.221, *p* = .036, *d* = .43), but less so when incongruent (imagining a reach but seeing a withdrawal, *M* = -.17, *SD* = .44, *t* = 2.0, *p* = .060, *d* = .39; imagining a withdrawal and seeing a reach, *M* = .13, *SD* = .43, *t* = 1.55, *p* = .133, *d* = .31), in line with an additive integration of expectations generated from the motion kinematics and prior imagery into perceptual judgments.

### Discussion

Experiment 2 tested whether imagery-induced predictive biases could also be observed in a perceptual judgment task that relies less on working memory or motor processes, by asking participants to compare the last seen step of the action sequence with a probe stimulus that was either identical, displaced further along the trajectory or not as far. The results fully replicated Experiment 1. They revealed, first, the classic predictive perceptual bias in the direction of motion (i.e., representational momentum effect, Freyd & Finke, 1984; Hubbard, 2005, 2018), by showing that probes displaced towards or away from the object were – erroneously – accepted more readily as “same” when the actor’s hand reached for and withdraw from the object, respectively, in line with a spontaneous filling in of the suddenly missing motion steps. Second, the results replicated the finding that imagery can induce similar predictive biases as actions that were indeed perceived, such that participants were more likely to mis-identify probes displaced towards the *imagined* trajectory, not only towards the expected one. Thus, when participants visualized a reach, they were more ready to erroneously accept as “same” probes shifted towards the object than probes shifted away from it. Conversely, when they imagined a withdrawal, they were more likely to judge probes shifted away from the object as “same” than those shifted towards it. Thus, as in Experiment 1, the observed biases in perceptual judgements reflected joint contributions of the expectations derived from the action’s real and imagined trajectories, in line with an integration of multiple expectations of a moving object’s behaviour (Hubbard, 1994, 2005).

These results rule out that the observed predictive biases in Experiment 1 were due to changes in action’s representation emerging in later working memory or motor control stages (e.g., spatial motor maps that guide hands to touch screen locations in Experiment 1). Instead, they point towards a contribution of imagery on primarily visuo-perceptual components of an action’s perceptual representation (i.e., filling in, iconic memory), similar as expectations derived from actual stimulus motion. Moreover, they rule out that the biases simply reflect motoric biases, as participants did not have to make any motor responses towards the target, but simply pressed a space bar. Finally, the probe judgment task is relatively rapid and non-transparent for participants, and the forward displacements it measures persist, at least partially, even when participants warned about them and given accurate feedback about their performance (Courtney & Hubbard, 2008; Freyd, 1987; Kelly & Freyd, 1987). The results are therefore unlikely to reflect demand or other higher-level biases.

## Experiment 3

Experiments 1 and 2 showed that prior mental imagery of action induces similar perceptual expectations as those generated during the observation of the actions themselves, so that both jointly determine the actions’ perceptual representation. Experiment 3 now tests whether imagery is not only integrated with expectations derived from the observed motion (i.e., representational momentum), but with more sophisticated expectations of how others typically act in given situations (Bach et al., 2014; Bach & Schenke, 2017), which do not simply bias perception towards or away from goal objects, but towards the particular trajectory people will take to achieve their goals. A hallmark of human social perception is the understanding that other people – and intentional agents in general – act efficiently, that is, that they try to minimize energy expenditure to achieve their goals within the given environmental constraints (e.g., “the intentional stance”, Baillargeon et al., 2016; Baker et al., 2009; Csibra & Gergely, 2007; Dennett, 1987; Gergely & Csibra, 2003; Scholl & Gao, 2013). An actor that is aware of the relevant features of the environment would therefore always take the straightest path towards a goal but would expend additional energy when they need to circumvent obstacles that are in the way.

The understanding that efficiency governs other people’s behaviour is acquired early in childhood and has been argued to provide the foundation of more sophisticated mentalizing skills, such as a prototypical theory of mind (as deviations from efficient actions signal that the actor has different goals, or considers different environmental constraints, as oneself; Csibra & Gergely, 2007; Gergely & Csibra, 2003). Importantly, a recent series of studies has shown that this understanding is not just represented in terms of abstract, symbolic knowledge of how others usually act, but is at least partially perceptually represented, so that it can be measured in representational momentum-like tasks (e.g., Hudson, Bach, et al., 2018; McDonough et al., 2019; McDonough & Bach, accepted). These studies show that people’s reports of others’ actions are biased not only towards their implied goal, or the future steps in their trajectory, but towards an ideal, energy-efficient trajectory with which these goals can be achieved. Thus, when watching a straight reach towards an obstacle, perceptual reports are subtly biased upwards, in line with the expectation that the actor would attempt to circumvent the obstacle. Conversely, when people observe an unnecessarily high, arched reach in the absence of an obstacle, perceptual reports of the hands’ last seen locations are “corrected” towards a more efficient straighter path. These biases are found as long as the actor is an intentional agent (instead of a mind-less ball, McDonough et al., 2019). Moreover, while the biases are larger when participants explicitly evaluate the most efficient trajectory before the action starts, or judge the presence of obstacles, they are still present even without any instruction (Hudson, Bach, et al., 2018; McDonough & Bach, accepted).

Experiment 3 tests whether imagery-induced action expectations are integrated with these expectations towards efficient action. In each trial, people saw a hand poised to reach a goal object on the other side of a table, with the path either unobstructed or obstructed by an obstacle, so that either a high arched reach (when an obstacle was present) or a flatter straight reach (when the path was clear) were expected. As before, a colour cue prompted participants to either imagine a straight low reach or a high arched reach, independently of whether an obstacle was present or not. In this way, the imagined action could be expected (e.g., imagining an arched reach when an obstacle was obstructing the path) or unexpected (e.g., imagining an arched reach when there was no obstacle to overcome). Participants then saw the hand begin to either make a straight or arched reach and they reported the hands’ last seen location after it had suddenly disappeared.

In contrast to Experiments 1 and 2, the current experiment does not measure whether imagery is integrated with expectations derived from prior motion, but with higher-order expectations about how intentional agents act within given environmental constraints. If imagery induces perceptual expectations about the action’s expected path, then expectations from imagery and efficient action should jointly affect perceptual reports, biasing perceptual reports not only towards the action continuations that are most efficient in the current environment, but also towards the imagined one. Reaches should then be reported to have disappeared higher not only when an obstacle was present but also when participants had imagined a high arched reach before the action commenced. In contrast, they should be reported lower in the absence of an obstacle and when people had just imagined a low straight reach.

### Method

#### Participants

Thirty-five participants took part in the experiment (mean age = 22 years, SD = 4.0, 21 females). All participants had normal/corrected-to-normal vision, gave informed consent, and were recruited from the University of Aberdeen for course credit or the Prolific community for payment. The study was approved by the University of Aberdeen’s ethics board, in line with the ESRC and the Declaration of Helsinki. The total number of 35 participants tested was preregistered, accounting for often high exclusion rates in online studies, with the goal to have at least 21 participants in the final sample, for 90% power to detect effects in the range of *d* = .76 (derived from prior work with this experimental paradigm and piloting). Six participants were excluded based on preregistered exclusion criteria (see Exclusion Criteria). The final sample size of 29 provides 90% power to detect effects of *d* > .62.

#### Preregistration

The study design and analysis plan was preregistered at aspredicted.org (https://aspredicted.org/8C1_MY6)^1^.

#### Apparatus and stimuli

Inquisit (Millisecond) software was used to present the experiment online via SONA and Prolific participant recruitment platforms.

Stimuli were presented at 50% of screen size and depicted an actor reaching for an object with either an efficient or inefficient trajectory, taken from prior studies (Hudson, McDonough, et al., 2018; McDonough et al., 2019; McDonough & Bach, accepted). They were derived from videos showing an arm that reaches from a resting position on the right of the screen towards a target object on the left (either an apple, a packet of crisps, a glue stick, or a stapler). The arm reached either straight for the target object (Straight/Efficient) or arched over an obstacle placed in the way (either an iPad, a table lamp or pencil holder; Arched/Efficient). Each video clip was converted into individual frames, and the first 22 from frame 1 (initial rest position) to 22 (mid-way through the action) were used as stimuli. For each efficient action, an inefficient action sequence was created by digitally removing the obstacles from the Arched/Efficient videos (Arched/Inefficient), or by inserting the obstructing objects into the Straight/Efficient videos, (Straight/Inefficient). The inefficient actions were therefore identical to the efficient actions in terms of movement kinematics and differed only by the presence/absence of the obstacle. Response stimuli were created by taking one frame from each action sequence and digitally removing the actor’s arm, so that only the objects and background remained. Presenting this frame immediately after the action sequence gave the impression of the hand disappearing from the scene, and participants indicated the last seen location of the tip of the index finger on this frame with a touch response on screen.

#### Procedure

To familiarize participants with the actions they would see and had to imagine, examples of each type of reach (straight with obstacle, straight without obstacle, arched with obstacle, arched without obstacle) were presented as videos in the instructions. Before the experiment started, participants completed a short training showing a random selection of trials from the main experiment. They were given the opportunity to repeat instructions and training if the task was unclear. Participants then completed a total of 96 trials, across two blocks, in which each type of trial was presented three times (2 reaches x 2 efficiencies x 2 imagined reaches x 4 sequence lengths).

An example trial sequence can be seen in Figure 5. At the start of each trial, participants saw the first frame of the action sequence as a static image with either a blue border or a purple border around it. The border thickness size was set at 5% of the width and height of the screen (which varied between our online participants). When the border was blue, participants were instructed to imagine the on-screen hand reaching for the target object with a straight-forward trajectory, even if there was an obstacle in the way. When the border was purple, participants were instructed to imagine the hand reaching with an arched trajectory, even if there was no obstacle to overcome. Text underneath this image reminded participants of these instructions. As soon as they felt that their mental image was clear and vivid, they pressed the spacebar, and the action sequence began 1000ms after.

**Figure 5.**
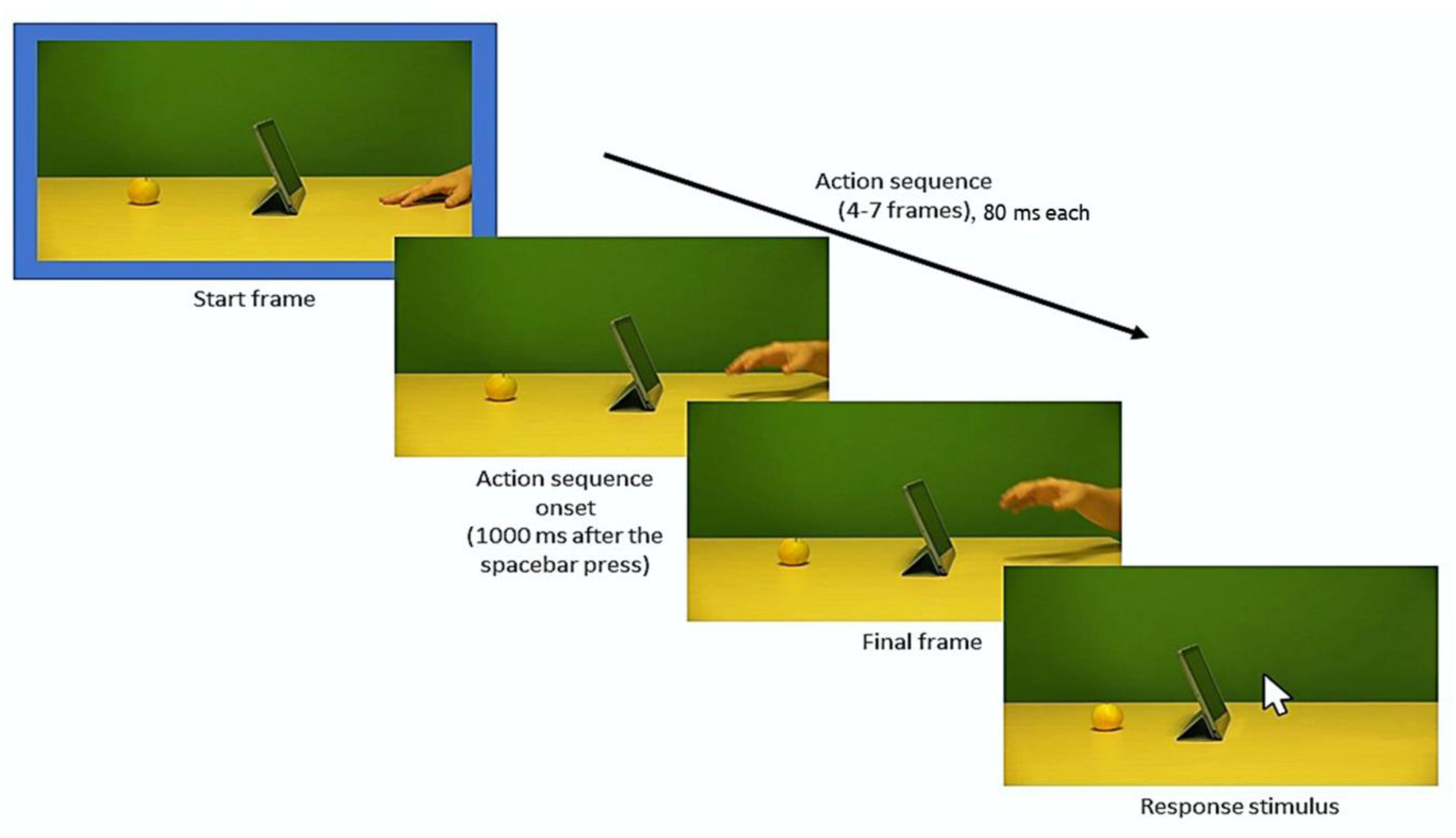
Illustration of the trial sequence in Experiment 3. Participants saw a hand in neutral position near an object, with a potential obstacle in between. Depending on the colour of the display frame they imagined the hand to either make a high arched reach or straight reach towards the goal, irrespective of the presence of an obstacle. When their imagery was vivid and clear, they pressed the space bar. This initiated the action, showing the hand making a high arched reach or a straight reach. After the hand disappeared, participants reported the index finger’s last seen location with the mouse.

The type of reach presented (Straight/Efficient, Straight/Inefficient, Arched/Efficient, Arched/Inefficient) was independent from the imagined reach. Every third frame of the action sequence was presented for 80ms each, with a randomly selected sequence length of 4, 5, 6 or 7 frames (e.g., trials with a length of 7 frames showed frames 1-4-7-10-13-16-19). The mouse cursor was hidden while the sequence played to prevent people from tracing its trajectory.

The final frame was replaced with the response frame, which showed the same scene without the hand, creating the impression that the hand had simply disappeared. The mouse pointer re-appeared, and participants used it to click on the screen to the report the final position of the tip of the index finger. As soon as a response was registered, the next trial began.

After all trials were completed, participants were given open-ended questions that asked (1) about any reasons why their data should not be used (e.g., because they were distracted), (2) to make a guess about the hypotheses tested by the experiment, and (3) whether there was any other feedback they wanted to leave us.

#### Exclusion criteria

Data exclusions followed preregistered criteria. Individual trials were excluded if the response time between hand disappearance and mouse localisation response was faster than 200ms or more than 3SDs above the sample mean (1.9%, *M* =1144ms, *SD* =1560). We additionally excluded trials (0.7%) in which participants did not make use of the imagery interval and terminated it before imagery could take place (< 200ms) and when the localisation error was greater than 300 pixels. While these additional criteria were not preregistered, their addition does not affect the pattern of results.

As specified in the preregistration, participants were excluded if the mean distance between the real final coordinates and their responses exceeded 3SDs of the sample mean (mean =131px, *SD* =166, one exclusion) or if the correlation between the real final coordinates and participant responses across trials was smaller than .70 (X axis: median r =.913, SD = .292; Y axis: median r =.883, SD = .288, four exclusions) [Footnote ^2^]. Participants with less than 50% of trials remaining were also removed (one exclusion). Finally, participants were removed if they indicated any reasons in their feedback to suggest that their data should not be used, for example, if they were distracted/interrupted during the task (no exclusions).

### Results

Figure 6 shows participants’ average perceptual reports of the hands’ disappearance points, as well as the average real trajectories that were presented. As online participants’ screen equipment differed, real and reported screen coordinates were mapped onto a screen of 960 x 540, with the zero point in the middle of the screen. As in Experiment 1, analysis was conducted on participants’ localization error, derived by subtracting the real final screen coordinate of the tip of the index finger from participants’ selected screen coordinate on each trial. This provides a directional measure of how far, in pixels (px), participants’ responses were displaced along the X- and Y-axes. An accurate response would produce a value of 0 on both axes. On the Y-axis, positive and negative values denote upward and downward displacements, respectively. On the X-axis, positive values denote a rightward displacement away from the object and negative values a leftward displacement.

**Figure 6.**
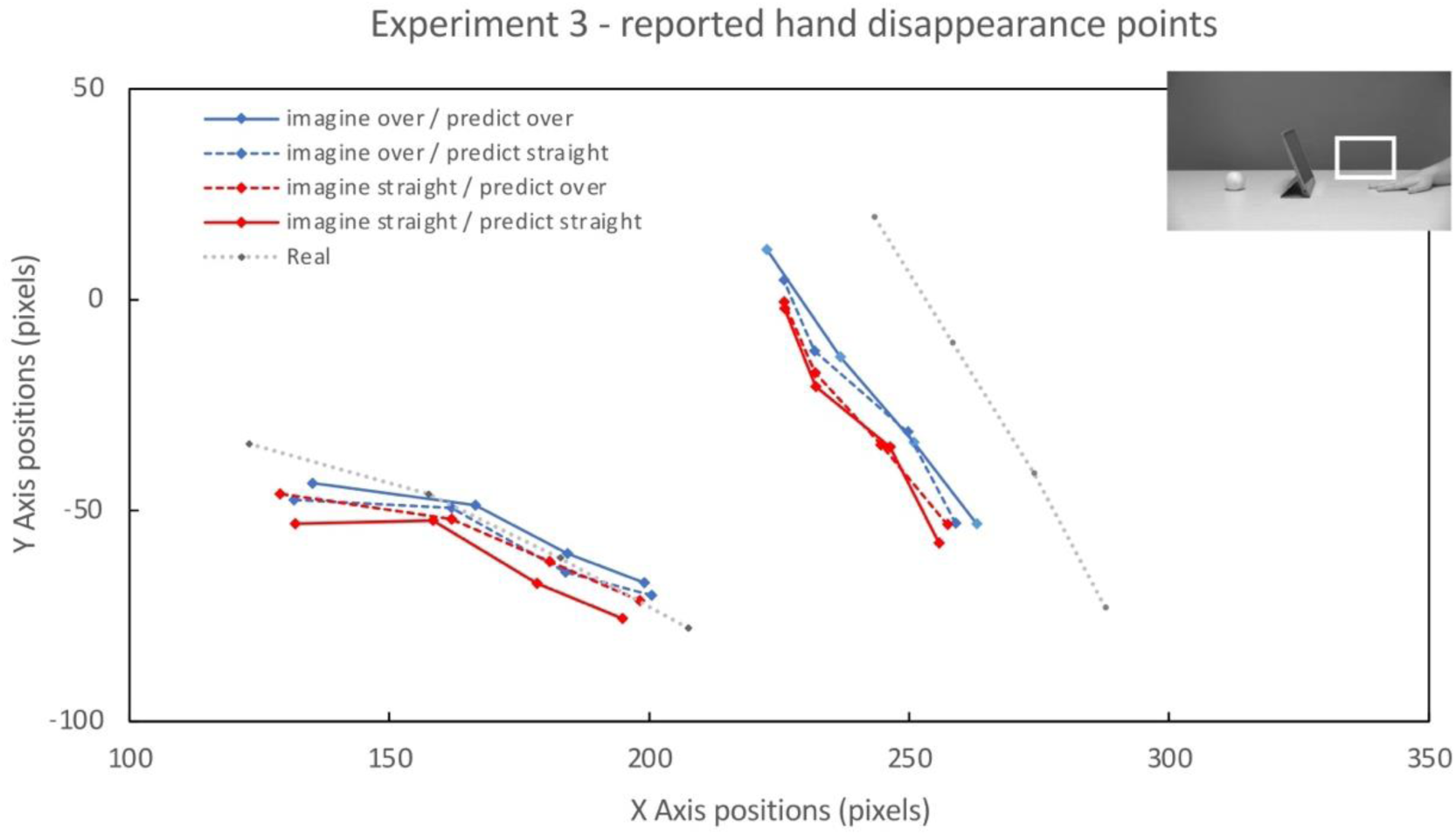
Absolute coordinates of reported and real hand disappearance points in Experiment 3, for both types of observed action (arched reach, straight reach), for each of the four disappearance points along their trajectories, depending on whether participants had imagined an action that matched the action predicted in the current environmental constraints (e.g., imagining an arched reach while an arched reach over an obstacle is expected; filled lines) or an action that did not match the environmental constraints (e.g., imagining an arched reach while the path is clear and a straight reach is expected; dotted lines). Grey dotted lines mark the real hand disappearance points. Coordinates are mapped onto a standardized screen of 960 x 540 pixels, where the origin (0,0) marks the centre of the screen and roughly corresponds to the obstacle location. Filled dots on each line mark the four possible disappearance points of the tip of the index finger along the reach trajectories. The rectangle in the inlay in the top right corner shows the stimulus section covered by the coordinate plot.

The difference scores were entered into a 2x2x2 ANOVA for the X and Y axis separately, with Observed Reach (straight vs arched), Imagined Reach (straight vs arched), and Expected Reach (straight vs arched) as repeated-measures factors [Footnote ^3^]. The main effects of Expected Reach and Imagined Reach on the Y axis will test our main hypothesis and indicate whether prior expectations derived from action efficiency and imagery jointly affect perceptual judgments. If perceptual judgments integrate both types of expectation, then they should be biased upwards not only when imagining an arched reach but particularly when such an arched reach is also expected due to an obstacle in the way. Similarly, perceptual judgments should be biased more strongly downwards not only when a straight reach was imagined, but also when such a reach is expected because the path to the goal object is obstacle-free.

#### Y axis

The analysis of perceptual errors on the Y Axis tests our hypotheses, that is, whether perceptual errors reflect joint contributions of the imagined trajectories and expectations about efficient actions given the current environmental constraints. Consistent with this prediction, the analysis revealed, first, a main effect of Expected Reach, *F*(1,28) = 7.11, *p* = .009, *ηp^2^*= .218, showing the contribution of expectations about the most efficient trajectory within the given the environmental constraints. Reported disappearance points were shifted upwards when the presence of an obstacle made an arched reach trajectory most likely (*M* = 0.8 px, *SD* = 10.4) and downwards when the path was clear and a straight reach was expected (*M* = -1.5 px, SD = 10.2). Importantly, the analysis also showed the predicted main effect of Imagined Reach, *F*(1,28) = 7.11, *p* = .013, *ηp^2^*= .203, revealing an independent contribution of prior imagery to perceptual judgments. Disappearance points were localised higher after arched reaches were imagined (*M* = 1.4 px, *SD* = 10.1) and lower after straight reaches were imagined (*M* = -2.1 px, *SD* = 11.7). As predicted, therefore, expectations from prior imagery were dynamically integrated with expectations of efficient action, so that the combined biases (Figure 6) upwards and downwards were largest when both influences were congruent (i.e., an arched reach is imagined and expected from the presence of an obstacle) and reduced when both were incongruent (i.e., an arched reach is imagined when a straight path towards the goal object was possible).

As we had no other predictions for the Y axis, all further ANOVA results should be treated as incidental, until they pass a threshold of *p* = .007 corrected for multiple comparisons in 2x2x2 ANOVA (Cramer et al., 2016). No tests met this threshold (all *F* < .896, *p* > .353), with the exception of a main effect of Observed Action, *F*(1,28) = 8.89, *p* < .006, *ηp^2^*= .241. Upwards- going arched reaches were mis-reported to have disappeared higher than they really did (*M* = 1.7 px, *SD* = 11.0) while straight reaches were reported to have disappeared lower (*M* = -2.3 px, *SD* = 10.9). While our study was not designed to reveal such influences, this is again consistent with the expectation that motions will continue on their previous path (i.e., representational momentum) measured in Experiments 1 and 2 and matches our prior research (Hudson, Bach, et al., 2018), which also revealed such effects.

#### X axis

We had no specific theoretical predictions for the perceptual errors on the X Axis. Therefore, all results should be treated as incidental, until they pass a threshold of *p* < .007, corrected for multiple comparisons in an ANOVA (Cramer et al., 2016). Two tests passed this threshold. A main effect of Observed Action, *F*(1,28) = 186.4, *p* < .001, *ηp^2^*= .869, showed that perceived disappearance points of straight reaches were shifted more strongly rightwards (*M* = 0.8 px, *SD* = 14.1) than of arched reaches (*M* = -24.2 px, *SD* = 16.3). This matches prior work with these stimuli (Hudson, McDonough, et al., 2018), and likely reflects a more rightwards-shifted centre of the hands for straight than arched reaches, which is known to affect localisation judgments (Coren et al., 1972). Interestingly, the analysis also revealed a main effect of Imagined Action, which passed the corrected threshold, *F*(1,28) = 13.0, *p* = .001, *η_p_ ^2^*= .317. Prior imagery of straight reaches did not only shift reports higher (see Y-axis results), but also more leftwards, towards the efficient straight trajectory towards the goal object (*M* = -13.6 px, *SD* = 13.8), relative to imagery of arched reaches (*M* = -9.9 px, *SD* = 15.5). While not a priori predicted, this is consistent with the idea that imagining high, arched reaches does not only bias motion expectations further upwards, but also less forward than straight reaches, which are imagined to head straight to the goal object.

### Discussion

Experiment 3 tested, in a preregistered procedure, whether imagery of actions induces perceptual expectations that are integrated with the efficiency expectations that govern the observation of others’ actions. We independently varied whether participants were asked to imagine a high, arched reach or a lower straight reach towards the goal object, and whether such reaches were expected, based on the presence or absence of obstacles in the scene. We found that expectations derived from imagery, and from the most likely actions within the given environmental constraints, jointly biased perceptual judgments. Thus, perceptual reports of hand disappearance were not only biased towards the most efficient action in a situation (i.e., upwards over an obstacle and downwards if the path is clear), but also towards the previously imagined kinematics (i.e., upwards for imagined arched than straight reaches). As a consequence, perceptual biases were most extreme when the different expectations were aligned (e.g., when people imagined an arched reach and an arched reach was expected) and reduced when both predicted opposing behaviours (e.g., when people imagined an arched reach while a straight one was expected).

These results show that prior imagery induces expectations of how motions will continue, which are integrated with other higher-order expectations about the observed action, so that they jointly bias perceptual judgments towards these expected next steps. These findings are in line with the proposal that imagery and prediction affect perception via shared pathways (e.g., Dijkstra et al., 2017, 2020; Moulton & Kosslyn, 2009), and that both influences are integrated into a joint representation about the observed motion’s next steps. What is unclear from the previous experiments is whether the influences from imagery and expectation converge only at a later stage when they become integrated with the visuospatial representation of the observed motion, or whether imagery can drive predictive processes directly, as would be expected if both make use of the same internal models and knowledge representations. Experiment 4 tests this idea.

## Experiment 4

Experiment 4 asks whether imagery is not only *integrated* with higher-order expectations of other people’s behaviour, but whether it can *drive* such expectations. If expectations and imagery make use of the same internal models, then imagery could be used to voluntarily shape one’s internal model of the world and to explore how these imagined situations would develop further. In the case of social perception, expectations about other people’s behaviour could then arise not (only) from the real world that people receive from their senses, but from the imagined world that they intentionally picture in their mind. Such an influence would make imagery highly useful for future planning. People could then imagine a situation, and predictive processes could take over to predict how others would behave within it. Such a strategy is common in everyday life when people imagine certain (potentially counterfactual) starting situations and then watch, in their minds eye, how they would unfold (Honeycutt et al., 1992; Honeycutt & Gotcher, 1991; Zagacki et al., 1992).

To test whether imagery can play such roles, we adapted the task of Experiment 3 to a different imagery instruction. Before each action started, the colour cue did not instruct participants to imagine different *actions* but different *situations*, that is, either that an obstacle interrupts the hand’s path to an object or that the path was clear, irrespective whether an obstacle is really present or not. This task therefore allows us to derive the independent contribution of the world model provided by the sensory input and by participants’ imagination in shaping expectations about others’ actions. Perceptual judgments of others’ action should then reflect both action expectations derived from the actual situation and be biased upwards when an obstacle was present or downwards when the path was clear, but show similar changes when these obstacles are merely imagined.

### Method

#### Participants

Forty-one participants took part in the experiment (mean age = 23.9 years, SD = 5.0, 21 female, 1 preferred not to say). All participants had normal/corrected-to-normal vision, gave informed consent, and were recruited from the University of Aberdeen for course credit or the Prolific community for payment. The study was approved by the University of Aberdeen’s ethics board, in line with the ESRC and the Declaration of Helsinki. Six participants were excluded based on preregistered exclusion criteria (see Exclusion Criteria). This number of exclusions was smaller than expected. Therefore, the final sample size of 35 surpassed our preregistered target sample size of at least 30, but results do not change if these additional five participants are removed.

#### Preregistration

The study design and analysis plan was preregistered at aspredicted.org (https://aspredicted.org/WVX_58X).

#### Apparatus, stimuli and procedure

The experiment was identical to Experiment 3, with the exception that the screen border’s colour now instructed participants about whether they had to imagine whether an obstacle (specifically, a table lamp) was present between the hand and goal object or not, and the number of possible obstacles to be presented was restricted to only this lamp stimulus so that participants had a clear mental image of which object to imagine. When the border was blue, participants were instructed to imagine that there was no object between hand and goal object, even if there actually was one in the way. When the border was purple, participants were instructed to imagine that a lamp was placed between hand and goal object, even if there currently was no such obstacle. As soon as they felt that their mental image was clear and vivid, they pressed the spacebar, and the action sequence began 1000ms after.

#### Exclusion criteria

Exclusion criteria followed our preregistered criteria and Experiment 3. Individual trials were excluded if the response times for the mouse localisation response after hand disappearance was faster than 200ms or more than 3SDs above the sample mean (1.2%, mean =703ms, *SD* =1560). We additionally excluded trials (0.2%) in which participants did not make use of the imagery interval and terminated it before any imagery could take place (< 200ms.) and when participants’ localisation error was greater than 300 pixels. While these additional criteria were not preregistered, their addition does not affect the pattern of results.

Participants were excluded if the mean distance between the real final coordinates and their responses exceeded 3SDs of the sample mean (mean =131px, *SD* =166, one exclusion) or if the correlation between the real final coordinates and participant responses were less than .70 across trials (X axis: median r =.913, SD = .292; Y axis: median r =.883, SD = .288, four exclusions). Participants with less than 50% trials remaining were also removed (no participants). Finally, participants were removed if they indicated any reasons in their feedback to suggest that their data should not be used, for example, if they were distracted/interrupted during the task (no participants).

### Results

Figure 7 shows participants localization of the hand disappearance points, as well as the real disappearance points. As online participants’ screen equipment differed between participants, real and reported screen coordinates were mapped onto a standardized resolution of 960 x 540 pixels. As in Experiment 1 and 3, analysis was conducted on participants’ localization error, derived by subtracting the real final screen coordinate of the tip of the index finger from participants’ selected screen coordinate on each trial.

**Figure 7.**
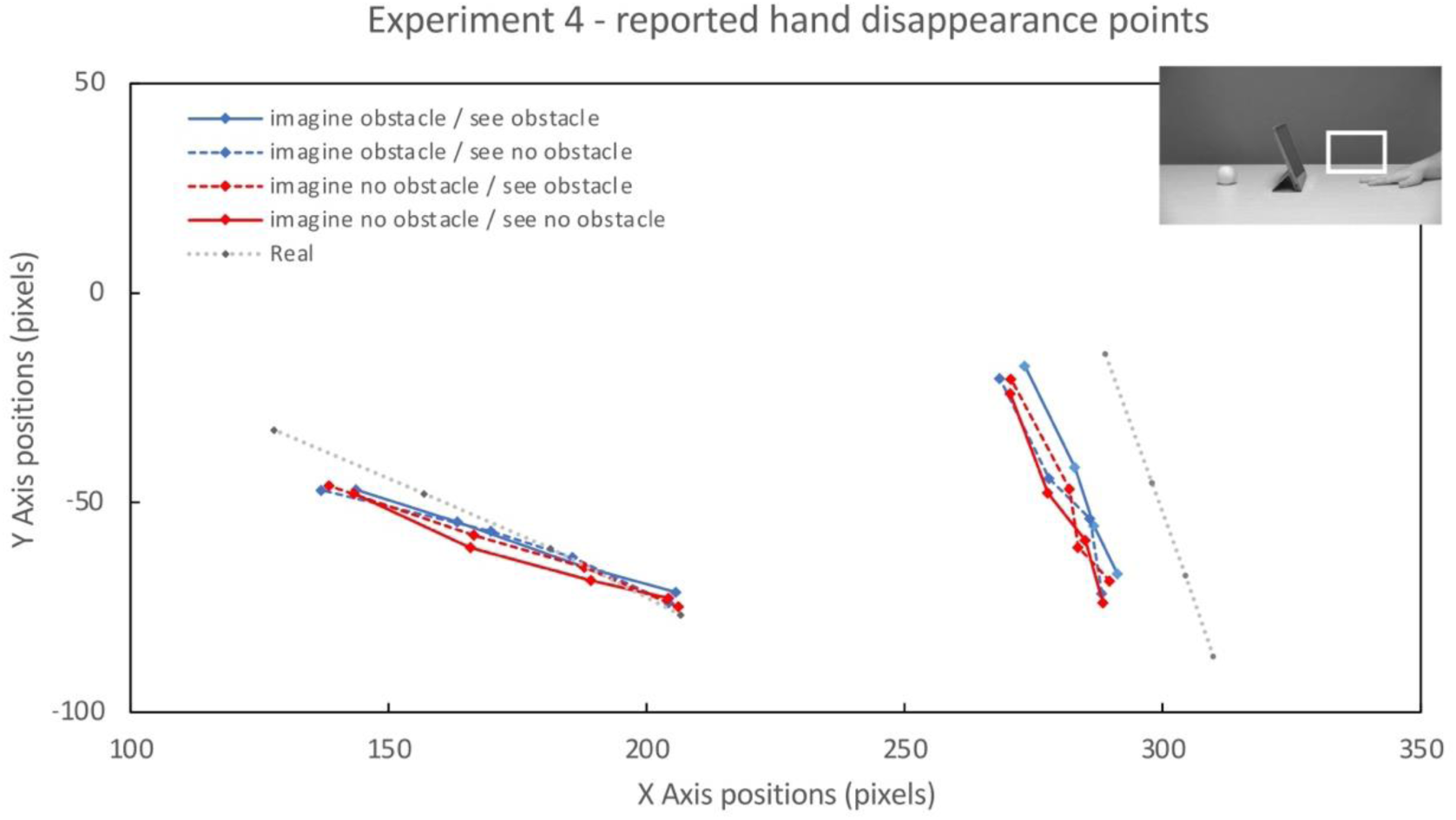
Absolute coordinates of reported and real hand disappearance points in Experiment 4, for both types of observed action (arched reach, straight reach), for each of the four disappearance points along their trajectories, depending on whether participants had imagined an obstacle in the way or the path being clear, and whether they saw an obstacle in the way or the path being clear. Grey dotted lines mark the real hand disappearance points. Coordinates are mapped onto a standardized screen of 960 x 540 pixels, where the origin (0,0) marks the centre of the screen and roughly corresponds to the obstacle location. Filled dots on each line mark the four possible disappearance points of the tip of the index finger along the reach trajectories. The rectangle in the inlay in the top right corner shows the stimulus section covered by the coordinate plot.

These difference scores were entered into a 2x2x2 ANOVA for the X and Y axis separately, with Observed Reach (straight vs arched), Real Scene (obstacle present, obstacle absent) and Imagined Scene (obstacle present, obstacle absent) as repeated-measures factors. Main effects of Observed Reach will reveal the extent to which prior action kinematics (arched vs straight reaches) bias perceptual reports in the prior motion direction (i.e., the representational momentum effect, Freyd & Finke, 1984; Hubbard, 2005), as also observed in Experiment 3. Main effects of Real Scene will reveal to what extent perceptual reports of reach disappearance points are biased towards the efficient trajectory given the real environmental constraints (upwards in the presence of an obstacle, straight when no obstacle is present).

Main effects of Imagined Scene will reveal how much perceptual reports are biased by the action that is expected in the imagined environment. The prediction is that perceptual judgements will be displaced upwards, towards the required arched trajectory, when an obstacle is imagined, and downwards, towards the straighter trajectory, when the obstacle is imagined to be absent and the path towards the goal is imagined to be clear.

#### Y axis

The analysis of perceptual displacements on the Y Axis tests our main hypotheses, that is, that perceptual reports will be displaced towards (1) the expected next steps in the sequence, (2) the trajectories expected in the given situation and (3) those expected for the imagined situation. The analysis revealed a main effect of Observed Reach, *F*(1,34) = 142.0, *p* < .001, *ηp^2^*= .807, replicating the overestimation of observed motion (i.e., representational momentum) towards the expected next steps in the sequence that was observed in Experiment 3. Upwards-going arched reaches were reported to have disappeared higher than they really did (*M* = 4.3 px, *SD* = 12.4) and straight reaches were reported to have disappeared slightly lower (*M* = -6.1 px, *SD* = 10.0). Importantly, the analysis also revealed the predicted main effect of Real Scene, *F*(1,34) = 12.4, *p* = .001, *ηp^2^*= .267, showing the previously demonstrated displacements towards the most efficient trajectory required in the current situation (Hudson et al., 2018; McDonough et al., 2019). Reaches were reported to have disappeared higher (*M* = 0.1 px, *SD* = 10.8) when an obstacle was present and an arched trajectory was expected than when there was no obstacle, and a straight reach was expected (*M* = -1.9 px, *SD* = 11.4). Finally, the analysis revealed the crucial main effect of Imagined Scene, *F*(1,34) = 14.9, *p* < .001, *ηp^2^*= .305. Just imagining an obstacle being present or absent changed perceptual reports similarly to a real object being present or absent. Imagined obstacles caused reports to be displaced more strongly upwards, towards the arched trajectory that would be required to circumvent this imagined obstacle (*M* = 0.6 px, *SD* = 11.3) than when no obstacle was imagined, and people could reach straight through this imagined scene (*M* = -2.4 px, *SD* = 11.1).

As we had no further predictions for the Y axis, all results further ANOVA results should be treated as incidental, until they pass a threshold of *p* = .007 corrected for multiple comparisons in explorative ANOVA (Cramer et al., 2016). No tests met this threshold (all *F* < .896, *p* > .353).

#### X axis

We had no specific predictions for perceptual displacements on the X Axis. Therefore, all results should be treated as incidental, until they pass a threshold of *p* = .007 corrected for multiple comparisons in an exploratory ANOVA (Cramer et al., 2016). The analysis only revealed a main effect of Observed Reach that would surpass this threshold, *F*(1,34) = 111.2, *p* < .001, *ηp^2^*= .766. Perceived disappearance points of arched reaches were displaced leftwards from the real disappearance points, towards the obstacle (*M* = -18.4 px, *SD* = 16.5) than of straight reaches (M = 7.6 px, *SD* = 14.7). Like in the original work with this experimental paradigm (Hudson, Bach, et al., 2018), this likely reflects the wider extent on the x axis for straight reaches, which leads to a right-displaced centre of gravity for the localisation judgments. It is interesting that, in contrast to Experiment 3, we did not find an influence of imagination on displacements on the X axis, *F*(1,34) = .182, *p* < .673, *ηp^2^*= .005, and that this difference between the experiments is robust, *F*(1,62) = 11.2, *p* < .001, *ηp^2^*= .153. While not a priori predicted, the effects of imagining obstacles therefore more closely mirror the effects of real obstacle presence, which also did not induce such biases on the X axis in either Experiment 3 or 4, than the effects of directly imaging arched reaches or straight reaches that head straight towards an obstacle.

### Discussion

Experiment 4 tested, in a preregistered procedure, whether imagery is not only integrated with prediction processes, but whether it can *drive* them, by measuring whether the measured perceptual biases integrate both action expectations induced by the actual presence of obstacles in the scenes, but also by obstacles that participants only imagined. The results replicated previous findings (Hudson, Bach, et al., 2018; Hudson, Nicholson, Ellis, et al., 2016; Hudson, Nicholson, Simpson, et al., 2016) that perceptual judgments of the hands’ disappearance points are biased towards the hand’s expected future trajectory (i.e., more upwards for arched than straight reaches), given the current environmental constraints (i.e., more upwards if an obstacle needs to be circumvented than when the path is clear). In addition, the experiment showed that simply imagining the presence and absence of obstacles induced similar perceptual (mis-)judgments. Thus, imagining an obstacle biased perceptual reports upwards, towards the trajectory required to circumvent this imagined obstacle, just as if an actual obstacle was present in the scenes. Conversely, high arched reaches were reported to have disappeared slightly lower when participants imagined a clear path towards the goal object that affords a straighter, less arched reach. Together, these data reveal that expectations from prior imagery are not only dynamically integrated with other expectations about how actions will continue, but that they can also drive these expectations, so that biases reflect not only the actions that are expected in the actual observed situation, but also the imagined one.

### Additional across-experiment analyses

#### Links to self-reported imagery strength

Previous studies have sometimes revealed links between behavioural measurements and individual differences in imagery strength (e.g., Lederman et al., 1990; Salge et al., 2021). In Experiments 1 and 2, we therefore correlated (1) participants’ perceptual biases in the motion direction (the representational momentum effect, Freyd & Finke, 1984; Hubbard, 2005) and (2) their perceptual biases towards the imagined action, with their scores on post-experiment questions about imagery strategy use, and responses on the VVIQ strength of imagery questionnaire. In Experiment 1, biases in the direction of motion reflected each participant’s contrast value for the main effect of Action, that is, how much their localisation errors were biased leftwards, in pixels, when observing reaches compared to withdrawals. Biases in the direction of imagery reflected the main effect of Imagery Prompt, that is, how much for leftwards perceptual errors were biased after imagining reaches compared to withdrawals. For Experiment 2, scores were derived analogously, capturing the interaction of Probe Direction and Action or Probe Direction and Imagery Prompt, capturing how much more likely participants were to accept leftwards-shifted probes as “same” compared to rightwards- shifted ones after seeing/imagining reaches compared to seeing/imagining withdrawals. To increase power for detecting any relationships, and because correlation scores are notoriously unreliable for low participant numbers (Hedge et al., 2018; Schönbrodt & Perugini, 2013), these tests were run over the pooled participants of Experiments 1 and 2, after standardizing results from each group by converting them into z-scores, from the touch screen and probe judgment tasks to make the two measurements comparable and to control for gross differences between the tasks and participant groups.

Overall, the post-experiment imagery scores were nicely interrelated. We found that responses on the VVIQ were positively related to participants’ post-experiment reports of their ability to control (*r* = .356, *p* = .009) and vividly experience (*r* = .398, *p* = .003) their imagery before each action, and to their reported use of visual (*r* = .330, *p* = .016) but not kinaesthetic (*r* = .023, *p* = .871) strategies in the task. Among the post-action scores, their reliance on visual imagery strategies was related to their experienced control (*r* = .422, *p* < .001) and vividness of their pre-action imagery (*r* = .363, *p* = .005). These interrelationships therefore provide some evidence that our questions measure aspects of imagery that overlap with the VVIQ questionnaire. Regarding the measured perceptual biases, we found that participants’ report of imagery use was broadly related to their tendency to overestimate the action trajectories they imagined towards the anticipated next steps (i.e., the representational momentum effect). Participants’ perceptual biases in the motion direction (the representational momentum effect, Freyd & Finke, 1984; Hubbard, 2005) correlated with their use of visual strategies (*r* = .350, *p* = .013). Moreover, there was some suggestive evidence for a correlation of representational momentum and participants’ scores on the VVIQ, but the correlation was not significant (*r* = .252, *p* = .077). While scores were still positive, participants’ overestimation of motion was however not robustly related to their use of kinaesthetic strategies (*r* = .200, *p* = .164), nor to the experienced control over their pre-action imagery (*r* = .194, *p* = .177) or its vividness (*r* = .114, *p* = .429). These data therefore provide some suggestive evidence of a link between the representational momentum effect and imagistic abilities, as reported before (Hubbard, 2006, 2017; Munger et al., 1999).

No reliable relationships were observed between subjective ratings of their imaginistic abilities and participants’ bias towards the imagined trajectory. The bias towards the imagined trajectory was neither robustly related to participants’ VVIQ scores (*r* = .032, *p* = .823), nor to their post-experiment reports on their ability to control imagery during the experiment (*r* = -.016, *p* = .912), nor its vividness (*r* = .015, *p* = .915). It was unrelated to the use of visual imagery strategies (*r* = .019, *p* = .898) but, interestingly, potentially weakly negatively related to the use of kinaesthetic imagery strategies (*r* = -.272, *p* = .056). Thus, while participants post-experiment imagery ratings were somewhat related to their general tendency to overestimate the observed actions towards their anticipated next steps, we did not observe any robust to relationships to imagery-induced localisation errors. Note though that even with the combined data of both experiments, our sample sizes are still not large enough to robustly reveal associations of robust behaviour tasks and individual difference measures (Hedge et al., 2018), and should therefore be interpreted with caution.

#### Links to guessing of the experimental hypotheses

A possibility is that some component of the measured perceptual biases in our experiments reflect demand effects, that is, that they emerge because participants guessed the experimental hypothesis during the experiment and shaped their responses accordingly.

Alternatively, participants’ ability to guess the hypothesis might be more generally related to their engagements with the experimental task. In Experiment 3 and 4, we therefore asked each participant after the experiment to make a (free text) guess of what hypothesis the experiment might have tested. Three raters independently evaluated each statement, blind with respect to which participant it came from, in how far the guess captured the idea of imagery-induced perceptual changes tested in the experiment. Each rater scored each response on a three-point scale that ranged from zero (the explanation does not match the experimental hypotheses, e.g., measuring reaction time), one (their explanation covers a general aspect of the experimental hypotheses, e.g., that imagery might broadly affect localisation accuracy), or two (their explanation captures the main part of the hypothesis, i.e., that imagining an arched reach will bias responses upwards more so than imagery of a straight reach). Across all 76 participants in the two experiments, the scores of the three raters were highly correlated, ranging from *r* = .69 to *r* =.78, suggesting adequate agreement. The median score of successful hypotheses guesses was *M* = .33, on the scale of 0 to 2, suggesting that most participants had no or little insight into the experimental hypotheses, however.

We then tested to what respect each participants’ average awareness score was related to measures of task compliance and their perceptual biases measured in the experiment. To increase power for detecting any relationships, and because correlation scores are notoriously unreliable for low participant numbers (Hedge et al., 2018; Schönbrodt & Perugini, 2013), these tests were again run over the pooled participants across both experiments.

We first checked whether awareness of the hypotheses was related to engagement with the experimental task, on the full set of participants before exclusion (as only participants with high correlations >.80 were considered for further analysis). We therefore correlated each participant’s average hypotheses guessing score to their (fisher-transformed) correlation score between their mouse localization responses and the hands’ real disappearance points. Indeed, both measures were correlated (*r* = .241, *p* = .031). Thus, perhaps not surprisingly, those participants who more fully engaged with the task and more faithfully localised the hand’s disappearance points were also more likely to successfully guess the experimental hypotheses after the experiment.

We then correlated the hypothesis guessing score of each participant that was considered for analysis in the two experiments to the three prediction measures of interest, measuring how much their perceptual judgments were distorted (1) in the direction of motion (more upwards for arched than straight reaches), (2) in the direction of the reach expected from the given environmental constraints (more upwards in the presence of an obstacle than empty space), and (3) in the direction of the imagined action (more upwards when imagining an arched reach/an obstacle than a straight reach/a clear path towards the goal). Note that only the third measure reflects the specific experimental hypothesis measured in the hypothesis ratings, while the others again provide some estimate for the general (baseline) link that can be expected because guessing the hypothesis is related to general task engagement.

No significant link between awareness of the hypotheses and any of the measures emerged in the tests of the non-standardized (raw) scores, for neither of the three relationships (motion extrapolation, *r* = .188; environmental constraints; *r* = .045; imagery, *r* = .172; all *p* > .138). When scores of the participants in each experiment were standardized via z-scores to control for gross differences between the two experiments, no relationships emerged either (motion extrapolation, *r* = .057; environmental constraints; *r* = .062; imagery, *r* = .202; all *p* > .109). Together, these findings provide little grounds to suspect a strong influence of demand effects. The relationships we observed are therefore more likely to be indicative of a general engagement with the experimental tasks that would increase hypothesis guesses and the measured perceptual biases towards expected and imagined trajectories.

## General Discussion

In four experiments, we tested whether mental imagery can be understood as the voluntary engagement of the same prediction processes that shape the perception of others’ behaviour. Participants imagined hand actions, such as reaches or withdrawals (Experiments 1 and 2), straight or arched reaches (Experiment 3), or environmental states in which such actions would be expected (presence or absence of obstacles, Experiment 4). We then measured how this prior imagery affects perception of the actual hand movements, by asking participants to report their last seen positions, either on a touch screen (Experiment 1), by comparing them to a probe stimulus (Experiment 2), or through mouse responses (Experiment 3 and 4).

All experiments revealed that imagery instructions induce motion expectations that distort perceptual reports towards the imagined paths, like other expectation-based influences on the observation of others’ behaviour. It is well-known that perceptual judgements of moving objects are predictively biased towards their expected future path. As a consequence, objects are reported to have travelled further along their trajectory than they really did (i.e., representational momentum, Freyd & Finke, 1984; Hubbard, 2005), with the specific extent and direction of these biases reflecting the combined expectations people have about how the motion will continue (for review, see Hubbard, 2005, 2018, 2020). Our findings replicated these biases, showing that moving hands were mis-perceived to have disappeared further along their path than they really did, both when reaching for or withdrawing from potential goal objects (Experiments 1 and 2) and when making straight or arched reaches (Experiments 3 and 4). Moreover, we also confirmed that these predictive biases were modulated by prior knowledge of how intentional agents navigate the environment, such that perceived hand disappearance points were biased towards a higher, more arched path when an obstacle needed to be overcome, and towards a straighter path when the path to the goal object was clear of any obstruction (Hudson, McDonough, et al., 2018; McDonough et al., 2019).

Importantly, all experiments showed that prior imagery of hand motions suffices to induce similar biases. In Experiments 1 and 2, perceptual mis-localisations towards the goal object were induced when participants had imagined a reach for the object, but biases away from the object were induced when people imagined a withdrawal. These imagery-induced biases were observed both when participants reported the hands’ perceived disappearance point on a touch screen (Experiment 1) and when they compared the hand’s location to a probe stimulus directly after (Experiment 2), indicating that these biases not only reflect changes to working memory or hand motor control (e.g., Kerzel, 2003; Müsseler et al., 2008). These findings are, to our knowledge, the first demonstration that imagery induces expectation-like biases during motion perception, which are dynamically integrated with other expectations towards the motions’ next steps, in line with the proposed integration of multiple sources of information during motion perception (see Hubbard, 1994; 2005).

Experiments 3 and 4 then tested whether the processes that form these mental images are integrated with, and can drive, higher-order expectations of how intentional agents typically act. In a preregistered procedure, Experiment 3 asked participants to imagine straight or arched reaches, in cases where such actions were expected given the environmental constraints (e.g., imagining an arched reach over an obstacle) or unexpected (imagining an arched reach when a straight path was clear). We found that, as in Experiments 1 and 2, perceptual judgements were biased towards the imagined trajectory: they were biased upwards when a high, arched reach was imagined, and downwards when a straight reach was imagined. Importantly, these biases were integrated with action expectations derived from the actual environment constraints, such that they were maximal when imagination matched these action expectations and minimal when they did not. Thus, imagining arched reaches biased judgments upwards more strongly when there really was an obstacle to overcome, but less so when a straight path was expected because the path was clear. Conversely, imagining a straight reach enhanced biases downwards when the path was clear, but less when an arched reach was expected due to an obstacle in the way.

Finally, Experiment 4 tested whether imagery is not only *integrated* into one’s predictions of a motion’s future path, but whether it can *drive* the formation of predictions, by acting directly on the internal models of the environment from which these expectations are derived. Participants were not asked to imagine certain actions, but different environmental constraints that would make specific actions more or less likely, such as whether an obstacle was obstructing (or not) a straight reach towards a goal. We then measured whether imagery of environmental constraints sufficed to induce motion expectations that agents in the scene would either reach around imagined obstacles or head straight towards goal objects for imagined straight paths. Indeed, in a preregistered procedure, we found that, irrespective of actual obstacle presence, simply imagining an obstacle caused participants to mis-locate observed hand trajectories slightly upwards, as expected for a hand that was avoiding the imagined obstacle. In contrast, imagining an unobstructed path distorted perceptual judgments downwards, towards a straight path towards the goal object. Experiment 4 therefore provides, to our knowledge, the first evidence that imagery is not only perception-like, or that it can induce predictive biases, but that it can act on the internal models of the environment from which others’ motion expectations are derived, showing that imagined environments can “stand in” for what is really perceived and drive the formation of further expectations, giving it a potential role in exploring counterfactual realities (Barlett & Brannon, 2006; Byrne, 2002; Redshaw & Suddendorf, 2020).

The findings we report here are in line with recent theoretical “predictive processing” proposals that liken imagery and predictive processes (Dijkstra et al., 2017, 2020; Grush, 2004; Moulton & Kosslyn, 2009). In these views, perception – social and otherwise – is seen as a top-down guided process of Bayesian hypothesis testing and revision. The expectations people have about the world and other people filter down the perceptual hierarchy and are translated into the perceptual input that would be received if these assumptions were correct, allowing them to be tested against the perceptual input and inform what is perceived (Clark, 2013; Den Ouden et al., 2012; Friston & Kiebel, 2009). Imagery, in such views, can be seen as an *intentional* mode of prediction-generation. It essentially inserts specific expectations – hypotheses about what one currently perceives – into this process, which therefore bias perceptual experience towards what is imagined just like other expectations. All findings – that imagery induces expectation-related changes to perceptual judgments (Experiments 1 to 3), that it provides the relevant informative context for expectation generation (Experiment 4), and that it biases perceptual representations of the observed motions – are in line with these proposals and argue for a fundamental sharing of resources between imagery, prediction, and ultimately the perceptual processes they act on (Dijkstra et al., 2017; Grush, 2004; Moulton & Kosslyn, 2009).

An important component of our findings is that the effects of imagery were measured through their effects on perceptual judgments, that is, how much imagery biased perceptual reports towards the induced expectations. Prior studies have provided evidence that imagery can act on – and become integrated with – perceptual experience, such that imagined stimuli provide additional evidence for perceptual judgements people make (e.g., Dijkstra et al., 2017). However, most of these studies used single, simple, abstract stationary stimuli, and measured whether – not how – these stimuli would be perceived (Farah, 1985; Pearson et al., 2008; Perky, 1910; Segal & Fusella, 1970). In contrast, the current studies showed that such imagery-perceptual interactions can be obtained for complex, naturalistic stimuli, and that they affect not only categorial (e.g., present/absent) judgments, but the kinematic representation of observed motions itself, inducing “illusory” changes to *how* they are visuospatially represented. Most importantly, the representational-momentum-tasks we used here measure these changes not as the fusion of two concurrent percepts – such as an imagined reach with an observed withdrawal – but in terms of the integration of multiple expectations people have about the moving stimulus’ *future* course, which then biases perceptual judgments towards this anticipated path (e.g., Hubbard, 1994; 2005).

At what level of processing do the effects emerge? While representational momentum had initially been conceptualized as a (working) memory-like effect (Freyd & Finke, 1984a, 1984b), it has since been demonstrated that it peaks rapidly after stimulus presentation (e.g., 260 to 500ms), and that it is related to processes in low-level perceptual regions such motion sensitive area MT (Senior et al., 2002). It is therefore likely that the biases either reflect “online” changes during motion observation (see Hogendoorn, 2020, for a review of similar findings from the Flash lag effect) or in the brief iconic/visual memory period directly after motion offset, which is relevant for establishing conscious representation of what was observed (Crick & Koch, 1990). As such, representational momentum may reflect mechanisms that play a role in resolving the considerable uncertainty during motion perception (i.e., motion blurring, Hammett, 1997) or which “fill in” the next steps of a motion sequence after a sudden offset (Ekman et al., 2017). Representational momentum, and representational momentum-like effects, are therefore increasingly understood as a “motion illusion” (Hubbard, 2005) and are likened other Bayesian-like integration phenomena during perceptual processing, such as the Flash lag effect (Hogendoorn, 2020; Nijhawan, 2002). The independence of the biases from response modality observed here across experiments (probe judgments, touchscreen judgments, mouse localisations) is consistent with such a lower-level locus. It is also supported by the finding that the effect can be observed in perceptual (probe judgment) tasks, directly after motion offset (250ms gap, Experiment 2) where working memory influences or motor biases are minimal (Freyd & Finke, 1984b; Hubbard, 2005, 2020; Hudson, Bach, et al., 2018; Hudson, Nicholson, Ellis, et al., 2016; Hudson, Nicholson, Simpson, et al., 2016), and by the dynamic integration of multiple imagery- and expectation-related sources into a joint perceptual representation found across all experiments here.

Moreover, the imagery-induced and other perceptual biases measured in our task were largely unrelated to participants’ guessing of the experimental hypotheses. Together with previous findings showing that representational momentum tasks are (at least partially) resistant against feedback manipulations (Courtney & Hubbard, 2008; Ruppel et al., 2009), this argues against a contribution of higher-level demand effects (i.e., Firestone & Scholl, 2016), but is consistent with the notion that the measured biases emerge from a more fundamental perceptual mechanism.

One (so far untested) prediction of such an integration of perception, imagery and expectations is that the effect of each component should increase with its reliability or “precision” (Ernst & Banks, 2002; Yon & Frith, 2021). By manipulating, between trials, the precision of the observed motions (e.g., brightness, visibility, smoothness), of participants’ action expectations (e.g., the likelihood that an actor would avoid obstacles), or, indeed, of the mental images they form (e.g., through imagery instructions, for example, to imagine a specific reach vs. a general motion to the left), future studies could measure how information from the different sources is weighted against each other and combined. Such investigations may resolve, why imagery sometimes seems to have perceptual consequences (if the precision of the imagined sensory features is high) but often appears to remain abstract (if precision of the sensory features is low), and therefore shed some light on conflicting findings in imagery-perception interactions Farah, 1985; Freyd & Finke, 1984b; Ishai & Sagi, 1995, 1997b; Pearson et al., 2008).

A general worry for imagery manipulations like ours is that the obtained effects may not be due to the imagery itself, but due to incidental features of the imagery instructions (e.g., Anderson, 1978; Zatorre & Halpern, 2005). In particular, representational momentum has been reported to be affected by prior verbal labels given to the motions (e.g., Hubbard, 1994; Hudson et al., 2016). Participants may therefore have spontaneously recoded the colour imagery cues we gave them into such verbal labels, which then generated the observed biases. Note however that, first, that the colour imagery cues in Experiments 1 and 2 were deliberately chosen to make such mnemonic “self-talk” unnecessary, with easy to remember colour-imagery associations (e.g., green=reach, red=withdraw), and we verified that these associations themselves did not induce any biases. Second, the labels of the to-be-imagined actions (“reach”, “withdraw”) by themselves held no directional information, without being instantiated in a given visuospatial scene with (e.g., with hands on the right and goal objects on the left). It is hard to see how such labels could induce any directional perceptual biases without being translated into an imaginistic representation of the resulting action’s path, as we explicitly instructed participants to do. This requirement for a perceptual instantiation is particularly pronounced in Experiment 4, where participants were only ever asked to imagine the presence or absence of a specific object (a lamp) between hand and goal object. Without some imaginistic integration with the scene context, it is hard to see how the labels “lamp” or “no lamp” could induce expectations towards a higher or lower reach, respectively. Finally, note that in prior studies on labelling on representational momentum, the effects were typically an order of magnitude smaller than those reported here, even with identical stimuli and measurement methods, and when the verbal labels were explicitly given and task relevant. For example, asking participants to verbalise (into a microphone) whether the actor should “take” or “leave” the object only induced biases of *ηp^2^* = .13, compared to *ηp^2^*=.42 in Experiment 2. In Experiment 1, imagery induced biases of *ηp^2^* = .65, but playing these verbal labels through the loudspeaker only induced effects of *ηp^2^*= .54, even if participants knew the labels accurately (75%) predicted the forthcoming actions (*ηp^2^*=.16 when they did not, Hudson et al., 2017). Last, in Experiment 3, imagery induced perceptual biases with *ηp^2^*= .20. In contrast, asking participants to mentally spell out the required reach (“straight”, “over”) only induced a non-significant effect of *ηp^2^*= .07, relative to asking participants to make verbally uninformative Yes/No judgments about obstacle presence (McDonough & Bach, accepted; see also Hudson, McDonough & Bach, 2018). Together, therefore, we see little grounds for a verbal mediation of our effects. Instead, we suspect that many previously reported effects of verbal labels occurred precisely because the given labels induced spontaneous imagery of motion within the given visuospatial context.

It is interesting to note that the motion extrapolation processes measured here have been shown before to be subject to other top-down influences, such as expectations derived from the physical forces acting upon the moving objects (e.g., friction, gravity; for reviews, see Hubbard, 2005, 2018b, 2020), one’s own actions (e.g., Jordan et al., 2002), or, in the domain of social perception, from the goals attributed to the moving actor (Hudson, Bach, et al., 2018; Hudson, Nicholson, Ellis, et al., 2016; Hudson, Nicholson, Simpson, et al., 2016) and the environmental constraints they have to navigate (Hudson, McDonough, et al., 2018; McDonough et al., 2019, see Experiments 3 and 4). Our data suggest that these – and other expectation-related effects in the literature – share a common basis with imagery, or, in fact, that expectation-related changes are mediated through spontaneous, imagery-like processes.

A fruitful avenue for future research would therefore be to test whether situations that induce predictive changes to perception generally do so because they elicit spontaneous mental images, which are then integrated with what is perceived (e.g., Moulton & Kosslyn, 2009; Neisser, 1976). One might then speculate for a fundamental identity of imagery and prediction processes; not only that imagery may rely on predictive internal models, but that prediction itself induces spontaneous imagery-like experiences.

Such an identity of prediction and imagery would go some way to explaining imagery’s fundamental role across many skills, from spatial navigation and reading comprehension to creativity, engineering, mathematics, moral decision-making (e.g., Blajenkova et al., 2006) and perspective taking (Ward et al., 2019, 2020, 2022). This role would then not only emerge from imagery’s ability to voluntarily generate counterfactual realities and visualise the perceptual input they would generate. It would also lie in the possibility for these imagined realities to interface with the predictive processes that instantiate the broader knowledge one has about how the physical and social reality most likely behaves (Hafri et al., 2022; for a similar argument in perspective taking, see Ward et al., 2019). If imagery can, as seen in Experiment 4, engage such predictive knowledge, it would provide us with perceptually instantiated information about how such starting situations would develop further, given one’s knowledge of the physical and social world. It would therefore enable imagery to support a broad range of skills – action planning, counterfactual thinking, moral reasoning, and theory of mind – which all require a targeted engagement of our world (and social) knowledge so that we can work out how a particular starting state would develop further, and which particular perceptual input it would generate.

## Conclusions

It has long been argued that imagery is a key tool for the anticipation of the relevant future (Dijkstra et al., 2017, 2020; Grush, 2004; Moulton & Kosslyn, 2009; Neisser, 1976), particularly when anticipating interactions with others (Honeycutt et al., 1992; Honeycutt & Gotcher, 1991; for reviews, see Honeycutt & Ford, 2001; Honeycutt & Gotcher, 1991). The present study shows that imagery of another’s action produces similar perceptual distortions as the prior expectations people have about how others’ actions will develop within a given situation. It therefore connects imagery to prediction processes in perceptual experience, specifically those in social perception, where the anticipation of others’ behaviour represents a fundamental channel for both understanding and coordinating social interactions. They therefore support the idea that imagery constitutes an offline, intentional activation of the brain’s internal model, which helps simulate situations not (yet) encountered and feed into the expectations we have of the world.

## Constraints on Generality

It is important to delineate some limitation of the current study, following recent recommendations to include an explicit statement describing the generality of a study’s findings (Simons et al., 2017). We expect that our results will generalize to any sample of participants in a research lab, and Experiments 3 and 4 confirmed that the general methodology generalize to an online setting and web-based experiments (sampled from populations beyond the psychology students usually tested in psychological studies). It should be noted that the stimuli in all experiments showed right hands moving towards (or away) from goal objects on the left. However, the central measurement technique implemented in representational momentum-like tasks has been shown to be widely generalizable and produce robust results across motion directions and stimulus types (actions, objects, abstract motions) (for reviews, see Hubbard, 2005, 2018b, 2020). Given the independence of the observed biases from the different sets of stimuli used here (e.g., reaches and withdrawals, straight and arched reaches, with different backgrounds and object contexts) and from the different response modalities used across experiments (probe judgments, touchscreen judgments, mouse localisations), we have no reason to believe that the present results depend on the characteristics of the response type, materials, or context. It would be important for future research to test the extent to which these results generalize across participants at different ages, whether they generalize to the developmental age or if such biases in perception may be related to evolutionary changes across species.

## Transparency and openness

All data sets and program codes used in this study are cited in the text and listed in the reference section. The data used for the analyses is available upon publication in an open repository here (https://osf.io/d9abp/). The study design and analysis plan of Experiment 3 (https://aspredicted.org/8C1_MY6) and Experiment 4 (https://aspredicted.org/WVX_58X) was preregistered at aspredicted.org; the preregistration link is provided for each study in the relative method section, and deviations from preregistration are made explicit. The present work includes replications of findings between experiments, as well as of previous findings reported in the literature.

## Author Note

The work was funded by Leverhulme Trust grant RPG-2019-248 to PB, and a PhD studentship awarded to Eleonora Parrotta from the Universities of Plymouth and Aberdeen. Below, we list the main authors’ contribution, in accordance with the CRediT taxonomy: Eleonora Parrotta: Conceptualization, Methodology, Software, Formal Analysis, Investigation, Writing, Visualization. Katrina McDonough: Conceptualization, Methodology, Software, Formal Analysis, Investigation, Writing, Visualization. Patric Bach: Conceptualization, Methodology, Formal Analysis, Writing, Visualization, Resources, Supervision, Project Administration, Funding Acquisition. The data reported in this manuscript have been presented at the UNSW Virtual workshop on Expectation, Perception and Cognition 2020; the Experimental Psychology Society meeting 2021, and the annual meeting of the Experimental Psychology Society UK in Stirling, Scotland in 2020.

## Footnotes

1. Our study design is based on prior work on Representational Momentum (e.g., Hubbard, 2005; Hubbard & Bharucha, 1988). There, representational momentum is often measured in terms of M and O displacements, where M-displacement indexes biases in the direction of motion (e.g., on the horizontal axis) and O-displacement denotes biases orthogonal to it (e.g., the vertical axis). Here, the main effect of Action on the X axis indexes the sum of how much further towards the object participants localise the hand’s disappearance point when seeing reaches and how much further away from the object they localise it for withdrawals. It therefore corresponds to the summed M-displacement induced by both action types. Analogously, the main effect of Imagery Prompt indexes how much further towards the object participants localise hand disappearance points when *imagining* reaches compared to withdrawals. It therefore roughly corresponds to the summed M-displacement induced by imagining reaches and withdrawals. For a separate estimation of the M-displacement in each condition, see the weighted means analysis in Experiment 2.
2. Note that, compared to Experiment 1, a slightly relaxed minimum correlation criterion of .70 was chosen (and preregistered) for Experiment 3 and 4, after piloting with this stimulus set with online samples revealed that such a relaxed criterion would substantially reduce participant exclusion without a major drop in effect sizes.
3. Please note that, for reasons of ease of understanding and consistency with the other experiments in the manuscript, the ANOVA model was coded differently than described in the preregistration. This does not affect the results nor the interpretation, with the only difference being that the crucial main effect of Expected Action here would emerge as an interaction of Imagined Action and Scene Congruency in the preregistered analysis, with equal statistical values.

## Notes

### Competing Interest Statement

The authors have declared no competing interest.

## References

Anderson, J. R. (1978). Arguments concerning representations for mental imagery. Psychological Review, 85(4), 249.

Bach, P., Nicholson, T., & Hudson, M. (2014). The affordance-matching hypothesis: How objects guide action understanding and prediction. Frontiers in Human Neuroscience, 8, 254.

Bach, P., & Schenke, K. C. (2017). Predictive social perception: Towards a unifying framework from action observation to person knowledge. Social and Personality Psychology Compass, 11(7), e12312.

Baillargeon, R., Scott, R. M., & Bian, L. (2016). Psychological reasoning in infancy. Annual Review of Psychology, 67(1), 159–186.

Baker, C. L., Saxe, R., & Tenenbaum, J. B. (2009). Action understanding as inverse planning. Cognition, 113(3), 329–349.

Barlett, C. P., & Brannon, L. A. (2006). “If only…”: The role of visual imagery in counterfactual thinking. Imagination, Cognition and Personality, 26(1), 87–100.

Bergmann, J., Genç, E., Kohler, A., Singer, W., & Pearson, J. (2016). Smaller primary visual cortex is associated with stronger, but less precise mental imagery. Cerebral Cortex, 26(9), 3838–3850.

Blajenkova, O., Kozhevnikov, M., & Motes, M. A. (2006). Object and spatial imagery: Distinctions between members of different professions. Cognitive Processing, 7, 20– 21.

Brandt, S. A., & Stark, L. W. (1997). Spontaneous eye movements during visual imagery reflect the content of the visual scene. Journal of Cognitive Neuroscience, 9(1), 27– 38.

Butter, C. M., Kosslyn, S., Mijovic-Prelec, D., & Riffle, A. (1997). Field-specific deficits in visual imagery following hemianopia due to unilateral occipital infarcts. Brain: A Journal of Neurology, 120(2), 217–228.

Byrne, R. M. (2002). Mental models and counterfactual thoughts about what might have been. Trends in Cognitive Sciences, 6(10), 426–431.

Campos, A., & Pérez-Fabello, M. J. (2009). Psychometric quality of a revised version Vividness of Visual Imagery Questionnaire. Perceptual and Motor Skills, 108(3), 798–802. https://doi.org/10.2466/PMS.108.3.798-802

Chang, S., Lewis, D. E., & Pearson, J. (2013). The functional effects of color perception and color imagery. Journal of Vision, 13(10), 4–4.

Chatterjee, A., & Southwood, M. H. (1995). Cortical blindness and visual imagery. Neurology, 45(12), 2189–2195.

Chiou, R., Rich, A. N., Rogers, S., & Pearson, J. (2018). Exploring the functional nature of synaesthetic colour: Dissociations from colour perception and imagery. Cognition, 177, 107–121.

Clark, A. (2013). Whatever next? Predictive brains, situated agents, and the future of cognitive science. Behavioral and Brain Sciences, 36(3), 181–204.

Coren, S., & Hoenig, P. (1972). Effect of non-target stimuli upon length of voluntary saccades. Perceptual and Motor Skills, 34(2), 499–508.

Courtney, J. R., & Hubbard, T. L. (2008). Spatial memory and explicit knowledge: An effect of instruction on representational momentum. Quarterly Journal of Experimental Psychology, 61(12), 1778–1784.

Cramer, A. O., van Ravenzwaaij, D., Matzke, D., Steingroever, H., Wetzels, R., Grasman, R. P., Waldorp, L. J., & Wagenmakers, E.-J. (2016). Hidden multiplicity in exploratory multiway ANOVA: Prevalence and remedies. Psychonomic Bulletin & Review, 23(2), 640–647.

Craver-Lemley, C., & Arterberry, M. E. (2001). Imagery-induced interference on a visual detection task. Spatial Vision.

Craver-Lemley, C., Arterberry, M. E., & Reeves, A. (1997). Effects of imagery on vernier acuity under conditions of induced depth. Journal of Experimental Psychology: Human Perception and Performance, 23(1), 3.

Craver-Lemley, C., & Reeves, A. (1987). Visual imagery selectively reduces vernier acuity. Perception, 16(5), 599–614.

Crick, F., & Koch, C. (1990). Towards a neurobiological theory of consciousness. Seminars in the Neurosciences, 2, 263–275.

Crowder, R. G. (1989). Imagery for musical timbre. Journal of Experimental Psychology: Human Perception and Performance, 15, 472–478. https://doi.org/10.1037/0096-1523.15.3.472

Csibra, G., & Gergely, G. (2007). ‘Obsessed with goals’: Functions and mechanisms of teleological interpretation of actions in humans. Acta Psychologica, 124(1), 60–78.

Delamillieure, P., Doucet, G., Mazoyer, B., Turbelin, M.-R., Delcroix, N., Mellet, E., Zago, L., Crivello, F., Petit, L., & Tzourio-Mazoyer, N. (2010). The resting state questionnaire: An introspective questionnaire for evaluation of inner experience during the conscious resting state. Brain Research Bulletin, 81(6), 565–573.

Den Ouden, H. E., Kok, P., & De Lange, F. P. (2012). How prediction errors shape perception, attention, and motivation. Frontiers in Psychology, 3, 548.

Dennett, D. C. (1987). The intentional stance. MIT press.

Dijkstra, N., Ambrogioni, L., Vidaurre, D., & van Gerven, M. (2020). Neural dynamics of perceptual inference and its reversal during imagery. Elife, 9, e53588.

Dijkstra, N., Bosch, S. E., & van Gerven, M. A. (2019). Shared neural mechanisms of visual perception and imagery. Trends in Cognitive Sciences, 23(5), 423–434.

Dijkstra, N., Mazor, M., Kok, P., & Fleming, S. (2021). Mistaking imagination for reality: Congruent mental imagery leads to more liberal perceptual detection. Cognition, 212, 104719.

Dijkstra, N., Zeidman, P., Ondobaka, S., van Gerven, M. A., & Friston, K. (2017). Distinct top-down and bottom-up brain connectivity during visual perception and imagery. Scientific Reports, 7(1), 1–9.

Ekman, M., Kok, P., & de Lange, F. P. (2017). Time-compressed preplay of anticipated events in human primary visual cortex. Nature Communications, 8(1), 1–9.

Ernst, M. O., & Banks, M. S. (2002). Humans integrate visual and haptic information in a statistically optimal fashion. Nature, 415(6870), 429–433.

Farah, M. J. (1985). Psychophysical evidence for a shared representational medium for mental images and percepts. Journal of Experimental Psychology: General, 114(1), 91.

Faul, F., Erdfelder, E., Lang, A.-G., & Buchner, A. (2007). G* Power 3: A flexible statistical power analysis program for the social, behavioral, and biomedical sciences. Behavior Research Methods, 39(2), 175–191.

Finks, R. A., Pinker, S., & Farah, M. J. (1989). Reinterpreting visual patterns in mental imagery. Cognitive Science, 13(1), 51–78.

Firestone, C., & Scholl, B. J. (2016). Cognition does not affect perception: Evaluating the evidence for “top-down” effects. Behavioral and Brain Sciences, 39.

Freyd, J. J. (1987). Dynamic mental representations. Psychological Review, 94(4), 427.

Freyd, J. J., & Finke, R. A. (1984a). Facilitation of length discrimination using real and imaged context frames. The American Journal of Psychology, 323–341.

Freyd, J. J., & Finke, R. A. (1984b). Representational momentum. Journal of Experimental Psychology: Learning, Memory, and Cognition, 10(1), 126.

Freyd, J. J., & Jones, K. T. (1994). Representational momentum for a spiral path. Journal of Experimental Psychology: Learning, Memory, and Cognition, 20(4), 968.

Friston, K., & Kiebel, S. (2009). Cortical circuits for perceptual inference. Neural Networks, 22(8), 1093–1104.

Gallagher, R. M., Suddendorf, T., & Arnold, D. H. (2021). The implied motion aftereffect changes decisions, but not confidence. *Attention, Perception*, & Psychophysics, 83(8), 3047–3055.

Ganis, G., & Schendan, H. E. (2008). Visual mental imagery and perception produce opposite adaptation effects on early brain potentials. Neuroimage, 42(4), 1714–1727.

Ganis, G., & Schendan, H. E. (2013). Cognitive neuroscience of mental imagery: Methods and paradigms. Multisensory Imagery, 283–298.

Ganis, G., Thompson, W. L., & Kosslyn, S. M. (2004). Brain areas underlying visual mental imagery and visual perception: An fMRI study. Cognitive Brain Research, 20(2), 226–241.

Gergely, G., & Csibra, G. (2003). Teleological reasoning in infancy: The naıve theory of rational action. Trends in Cognitive Sciences, 7(7), 287–292.

Goss, S., Hall, C., Buckolz, E., & Fishburne, G. (1986). Imagery ability and the acquisition and retention of movements. Memory & Cognition, 14(6), 469–477.

Grush, R. (2004). The emulation theory of representation: Motor control, imagery, and perception. Behavioral and Brain Sciences, 27(3), 377–396.

Hafri, A., Boger, T., & Firestone, C. (2022). Melting ice with your mind: Representational momentum for physical states. Psychological Science, 33(5), 725–735.

Hammett, S. T. (1997). Motion blur and motion sharpening in the human visual system. Vision Research, 37(18), 2505–2510.

Hedge, C., Powell, G., & Sumner, P. (2018). The reliability paradox: Why robust cognitive tasks do not produce reliable individual differences. Behavior Research Methods, 50(3), 1166–1186. https://doi.org/10.3758/s13428-017-0935-1

Hogendoorn, H. (2020). Motion extrapolation in visual processing: Lessons from 25 years of flash-lag debate. Journal of Neuroscience, 40(30), 5698–5705.

Honeycutt, J. M., & Ford, S. G. (2001). Mental imagery and intrapersonal communication: A review of research on imagined interactions (IIs) and current developments. Annals of the International Communication Association, 25(1), 315–345.

Honeycutt, J. M., & Gotcher, J. M. (1991). Influence of imagined interactions on communicative outcomes: The case of forensic competition. Mental Imagery, 139–143.

Honeycutt, J. M., Zagacki, K. S., & Edwards, R. (1992). Imagined interaction, conversational sensitivity and communication competence. Imagination, Cognition and Personality, 12(2), 139–157.

Housner, L., & Hoffman, S. J. (1981). Imagery ability in recall of distance and location information. Journal of Motor Behavior, 13(3), 207–223.

Hubbard, T. L. (1993). The effect of context on visual representational momentum. Memory & Cognition, 21(1), 103–114.

Hubbard, T. L. (1994). Judged Displacement: A Modular Process? The American Journal of Psychology, 107(3), 359–373. https://doi.org/10.2307/1422879

Hubbard, T. L. (1995a). Cognitive representation of motion: Evidence for representational friction and gravity analogues. Journal of Experimental Psychology: Learning, Memory, & Cognition, 21, 1–14.

Hubbard, T. L. (1995b). Environmental invariants in the representation of motion: Implied dynamics and representational momentum, gravity, friction, and centripetal force. Psychonomic Bulletin & Review, 2, 322–338.

Hubbard, T. L. (2005). Representational momentum and related displacements in spatial memory: A review of the findings. Psychonomic Bulletin & Review, 12(5), 822–851.

Hubbard, T. L. (2006). Bridging the gap: Possible roles and contributions of representational momentum. Psicológica.

Hubbard, T. L. (2017). Toward a general theory of momentum-like effects. Behavioural Processes, 141, 50–66.

Hubbard, T. L. (2018a). Influences on representational momentum. Spatial Biases in Perception and Cognition, 121–138.

Hubbard, T. L. (2018b). Spatial biases in perception and cognition. Cambridge University Press.

Hubbard, T. L. (2020). Representational gravity: Empirical findings and theoretical implications. Psychonomic Bulletin & Review, 27(1), 36–55.

Hubbard, T. L., & Bharucha, J. J. (1988). Judged displacement in apparent vertical and horizontal motion. Perception & Psychophysics, 44(3), 211–221.

Hubbard, T. L., & Stoeckig, K. (1988). Musical imagery: Generation of tones and chords. Journal of Experimental Psychology: Learning, Memory, and Cognition, 14(4), 656.

Hudson, M., Bach, P., & Nicholson, T. (2018). You said you would! The predictability of other’s behavior from their intentions determines predictive biases in action perception. Journal of Experimental Psychology: Human Perception and Performance, 44(2), 320.

Hudson, M., McDonough, K. L., Edwards, R., & Bach, P. (2018). Perceptual teleology: Expectations of action efficiency bias social perception. Proceedings of the Royal Society B, 285(1884), 20180638.

Hudson, M., Nicholson, T., Ellis, R., & Bach, P. (2016). I see what you say: Prior knowledge of other’s goals automatically biases the perception of their actions. Cognition, 146, 245–250.

Hudson, M., Nicholson, T., Simpson, W. A., Ellis, R., & Bach, P. (2016). One step ahead: The perceived kinematics of others’ actions are biased toward expected goals. Journal of Experimental Psychology: General, 145(1), 1.

Isaac, A. R., & Marks, D. F. (1994). Individual differences in mental imagery experience: Developmental changes and specialization. British Journal of Psychology, 85(4), 479–500.

Ishai, A., & Sagi, D. (1995). Common mechanisms of visual imagery and perception. Science, 268(5218), 1772–1774.

Ishai, A., & Sagi, D. (1997a). Visual imagery: Effects of short-and long-term memory. Journal of Cognitive Neuroscience, 9(6), 734–742.

Ishai, A., & Sagi, D. (1997b). Visual imagery facilitates visual perception: Psychophysical evidence. Journal of Cognitive Neuroscience, 9(4), 476–489.

Jack, A. I., & Roepstorff, A. (2002). Introspection and cognitive brain mapping: From stimulus–response to script–report. Trends in Cognitive Sciences, 6(8), 333–339.

Jordan, J. S., Stork, S., Knuf, L., Kerzel, D., & Müsseler, J. (2002). Action planning affects spatial localization. Attention and Performance XIX: Common Mechanisms in Perception and Action, 158–176.

Kelly, M. H., & Freyd, J. J. (1987). Explorations of representational momentum. Cognitive Psychology, 19(3), 369–401.

Kerzel, D. (2003). Mental extrapolation of target position is strongest with weak motion signals and motor responses. Vision Research, 43(25), 2623–2635.

Kilner, J. M., Friston, K. J., & Frith, C. D. (2007). Predictive coding: An account of the mirror neuron system. Cognitive Processing, 8(3), 159–166.

Kobayashi, M., Takeda, M., Hattori, N., Fukunaga, M., Sasabe, T., Inoue, N., Nagai, Y., Sawada, T., Sadato, N., & Watanabe, Y. (2004). Functional imaging of gustatory perception and imagery:“top-down” processing of gustatory signals. Neuroimage, 23(4), 1271–1282.

Koenig-Robert, R., & Pearson, J. (2021). Why do imagery and perception look and feel so different? Philosophical Transactions of the Royal Society B, 376(1817), 20190703.

Kosslyn, S. M. (1973). Scanning visual images: Some structural implications. Perception & Psychophysics, 14(1), 90–94.

Kosslyn, S. M. (1980). Image and mind. Harvard University Press.

Kosslyn, S. M., Ball, T. M., & Reiser, B. J. (1978). Visual images preserve metric spatial information: Evidence from studies of image scanning. Journal of Experimental Psychology: Human Perception and Performance, 4(1), 47.

Kosslyn, S. M., & Thompson, W. L. (2003). When is early visual cortex activated during visual mental imagery? Psychological Bulletin, 129(5), 723.

Laeng, B., & Teodorescu, D.-S. (2002). Eye scanpaths during visual imagery reenact those of perception of the same visual scene. Cognitive Science, 26(2), 207–231.

Lederman, S. J., Klatzky, R. L., Chataway, C., & Summers, C. D. (1990). Visual mediation and the haptic recognition of two-dimensional pictures of common objects. Perception & Psychophysics, 47(1), 54–64.

Markov, N. T., & Kennedy, H. (2013). The importance of being hierarchical. Current Opinion in Neurobiology, 23(2), 187–194.

Marks, D. F. (1995). New directions for mental imagery research.

McDonough, K. L., Hudson, M., & Bach, P. (2019). Cues to intention bias action perception toward the most efficient trajectory. Scientific Reports, 9(1), 1–10.

McKelvie, S. J. (1995). Response to commentaries: The VVIQ and beyond. Journal of Mental Imagery, 19, 197–251.

Monaco, S., Malfatti, G., Culham, J. C., Cattaneo, L., & Turella, L. (2020). Decoding motor imagery and action planning in the early visual cortex: Overlapping but distinct neural mechanisms. Neuroimage, 218, 116981.

Mostert, P., Albers, A. M., Brinkman, L., Todorova, L., Kok, P., & De Lange, F. P. (2018). Eye movement-related confounds in neural decoding of visual working memory representations. Eneuro, 5(4).

Moulton, S. T., & Kosslyn, S. M. (2009). Imagining predictions: Mental imagery as mental emulation. Philosophical Transactions of the Royal Society B: Biological Sciences, 364(1521), 1273–1280.

Mulder, T. (2007). Motor imagery and action observation: Cognitive tools for rehabilitation. Journal of Neural Transmission, 114, 1265–1278.

Munger, M. P., Solberg, J. L., Horrocks, K. K., & Preston, A. S. (1999). Representational momentum for rotations in depth: Effects of shading and axis. Journal of Experimental Psychology: Learning, Memory, and Cognition, 25(1), 157.

Müsseler, J., Stork, S., & Kerzel, D. (2008). Localizing the onset of moving stimuli by pointing or relative judgment: Variations in the size of the Fröhlich effect. Vision Research, 48(4), 611–617.

Naselaris, T., Olman, C. A., Stansbury, D. E., Ugurbil, K., & Gallant, J. L. (2015). A voxel-wise encoding model for early visual areas decodes mental images of remembered scenes. Neuroimage, 105, 215–228.

Neisser, U. (1976). Cognition and reality: Principles and implications of cognitive psychology (pp. xiii, 230). W H Freeman/Times Books/ Henry Holt & Co.

Nijhawan, R. (2002). Neural delays, visual motion and the flash-lag effect. Trends in Cognitive Sciences, 6(9), 387–393.

Pearson, J. (2019). The human imagination: The cognitive neuroscience of visual mental imagery. Nature Reviews Neuroscience, 20(10), 624–634.

Pearson, J., Clifford, C. W., & Tong, F. (2008). The functional impact of mental imagery on conscious perception. Current Biology, 18(13), 982–986.

Pearson, J., & Kosslyn, S. M. (2015). The heterogeneity of mental representation: Ending the imagery debate. Proceedings of the National Academy of Sciences, 112(33), 10089– 10092.

Perky, C. W. (1910). An experimental study of imagination. The American Journal of Psychology, 21(3), 422–452.

Porro, C. A., Francescato, M. P., Cettolo, V., Diamond, M. E., Baraldi, P., Zuiani, C., Bazzocchi, M., & Di Prampero, P. E. (1996). Primary motor and sensory cortex activation during motor performance and motor imagery: A functional magnetic resonance imaging study. Journal of Neuroscience, 16(23), 7688–7698.

Pylyshyn, Z. W. (1973). What the mind’s eye tells the mind’s brain: A critique of mental imagery. Psychological Bulletin, 80(1), 1.

Pylyshyn, Z. W. (1981). The imagery debate: Analogue media versus tacit knowledge. Psychological Review, 88(1), 16.

Redshaw, J., & Suddendorf, T. (2020). Temporal junctures in the mind. Trends in Cognitive Sciences, 24(1), 52–64.

Reeves, A. (1981). Visual imagery lowers sensitivity to hue-varying, but not to luminance-varying, visual stimuli. Perception & Psychophysics, 29, 247–250.

Ruppel, S. E., Fleming, C. N., & Hubbard, T. L. (2009). Representational momentum is not (totally) impervious to error feedback. Canadian Journal of Experimental Psychology/Revue Canadienne de Psychologie Expérimentale, 63(1), 49.

Salge, J. H., Pollmann, S., & Reeder, R. R. (2021). Anomalous visual experience is linked to perceptual uncertainty and visual imagery vividness. Psychological Research, 85(5), 1848–1865.

Schendan, H. E., & Ganis, G. (2012). Electrophysiological potentials reveal cortical mechanisms for mental imagery, mental simulation, and grounded (embodied) cognition. Frontiers in Psychology, 3, 329.

Scholl, B. J., & Gao, T. (2013). Perceiving animacy and intentionality: Visual processing or higher-level judgment. Social Perception: Detection and Interpretation of Animacy, Agency, and Intention, 4629, 197–229.

Schönbrodt, F. D., & Perugini, M. (2013). At what sample size do correlations stabilize? Journal of Research in Personality, 47(5), 609–612.

Segal, S. J., & Fusella, V. (1970). Influence of imaged pictures and sounds on detection of visual and auditory signals. Journal of Experimental Psychology, 83(3p1), 458.

Senior, C., Ward, J., & David, A. S. (2002). Representational momentum and the brain: An investigation into the functional necessity of V5/MT. Visual Cognition, 9(1–2), 81– 92.

Shepard, R. N., & Metzler, J. (1971). Mental rotation of three-dimensional objects. Science, 171(3972), 701–703.

Simons, D. J., Shoda, Y., & Lindsay, D. S. (2017). Constraints on generality (COG): A proposed addition to all empirical papers. Perspectives on Psychological Science, 12(6), 1123–1128.

Tamir, D. I., & Thornton, M. A. (2018). Modeling the predictive social mind. Trends in Cognitive Sciences, 22(3), 201–212.

Thornton, I., & Hayes, A. (2004). Anticipating action in complex scenes. Visual Cognition, 11(2–3), 341–370. https://doi.org/10.1080/13506280344000374

Toussaint, L., Tahej, P.-K., Thibaut, J.-P., Possamai, C.-A., & Badets, A. (2013). On the link between action planning and motor imagery: A developmental study. Experimental Brain Research, 231, 331–339.

Vandenberghe, A., & Vannuscorps, G. (2022). Predictive extrapolation of observed body movements is tuned by knowledge of the body biomechanics. Journal of Experimental Psychology: Human Perception and Performance.

Vannuscorps, G., & Caramazza, A. (2016). The origin of the biomechanical bias in apparent body movement perception. Neuropsychologia, 89, 281–286.

Ward, E., Ganis, G., & Bach, P. (2019). Spontaneous vicarious perception of the content of another’s visual perspective. Current Biology, 29(5), 874–880.

Ward, E., Ganis, G., McDonough, K. L., & Bach, P. (2020). Perspective taking as virtual navigation? Perceptual simulation of what others see reflects their location in space but not their gaze. Cognition, 199, 104241.

Ward, E., Ganis, G., McDonough, K. L., & Bach, P. (2022). Is implicit Level-2 visual perspective-taking embodied? Spontaneous perceptual simulation of others’ perspectives is not impaired by motor restriction. Quarterly Journal of Experimental Psychology, 75(7), 1244–1258.

Wilson, M., Lancaster, J., & Emmorey, K. (2010). Representational momentum for the human body: Awkwardness matters, experience does not. Cognition, 116(2), 242– 250.

Winawer, J., Huk, A. C., & Boroditsky, L. (2010). A motion aftereffect from visual imagery of motion. Cognition, 114(2), 276–284.

Yon, D., & Frith, C. D. (2021). Precision and the Bayesian brain. Current Biology, 31(17), R1026–R1032.

Zagacki, K. S., Edwards, R., & Honeycutt, J. M. (1992). The role of mental imagery and emotion in imagined interaction. Communication Quarterly, 40(1), 56–68.

Zatorre, R. J., & Halpern, A. R. (2005). Mental concerts: Musical imagery and auditory cortex. Neuron, 47(1), 9–12.

Zatorre, R. J., Halpern, A. R., Perry, D. W., Meyer, E., & Evans, A. C. (1996). Hearing in the mind’s ear: A PET investigation of musical imagery and perception. Journal of Cognitive Neuroscience, 8(1), 29–46.

